# *De novo* aging-related megaloneurites: alteration of NADPH diaphorase positivity in the sacral spinal cord of the aged dog

**DOI:** 10.1101/483990

**Authors:** Yinhua Li, Yunge Jia, Wei Hou, Zichun Wei, Xiaoxin Wen, Yu Tian, Weijin Zhang, Lu Bai, Anchen Guo, Guanghui Du, Huibing Tan

**Affiliations:** Department of Anatomy, Jinzhou Medical University, Jinzhou, Liaoning 121001, China; Key Laboratory of Neurodegenerative Diseases of Liaoning Province, Jinzhou Medical University, Jinzhou, Liaoning, 121001,China; Laboratory of Biological Enzyme, Qinhuangdao Yier Bio-tech Co. LTD., Qinghuangdao, Hebei, 06000, China; Laboratory of Clinical Medicine Research, Beijing Tiantan Hospital, Capital Medical University, Beijing 100050, China; Department of Urology, Tongji Medical College Affiliated Tongji Hospital, Huazhong University of Science and Technology, Wuhan, Hubei 430030, China

**Keywords:** Aging, Lumbosacral spinal cord, NADPH diaphorase, megaloneurite, vasoactive intestinal peptide, Dog

## Abstract

The aging-related changes of NADPH-diaphorase (NADPH-d) in the spinal cord were studied in aged dogs. At all levels of the spinal cord examined, NADPH-d activities were present in neurons and fibers in the superficial dorsal horn, dorsal commissure and in neurons around the central canal. In addition, the sympathetic autonomic nucleus in the thoracic and rostral lumbar segments exhibited prominent NADPH-d cellular staining whereas the sacral parasympathetic nucleus (SPN) in the sacral segments was not well stained. Interestingly, we found abundant NADPH-d positive enlarged-diameter fibers termed megaloneurite, which characteristically occurred in the aged sacral segments, distributed in the dorsal gray commissure (DGC), lateral collateral pathway (LCP) the lateral fasciculi and the central canal compared with the cervical, thoracic and lumbar segments. The dense, abnormal NADPH-d megaloneurites occurred in extending from dorsal entry zone through lamina I along with the lateral boundary of the dorsal horn to the region of the SPN. These fibers were prominent in the S1-S3 segments but not in adjacent segments L5-L7 and Cx1 or in thoracolumbar segments and cervical segments. Double staining with GFAP, NeuN, CGRP, MAP2 and Iba1, NADPH-d megaloneurite colocalized with vasoactive intestinal peptide. Presumably, the megaloneurites may represent, in part, visceral afferent projections to the SPN and/or DGC. The NADPH-d megaloneurites in the aged sacral spinal cord indicated some anomalous changes in the neurites, which might account for a disturbance in the aging pathway of the autonomic and sensory nerve in the pelvic visceral organs.

## Introduction

The caudal lumbar and sacral spinal cord is important for controlling the function of the large intestine, pelvic muscles and the urogenital organs (Berkley et al., 1993; Jobling et al., 2010; Cruz et al., 2017). The sacral spinal cord is more specifically related to the intestine, bladder and sexual dysfunction (Cohen et al., 1991; Ogiwara and Morota, 2014; Barbe et al., 2018). In the dorsal gray commissure (DGC) in the S1-S3 segments of the cat spinal cord, this region receives the terminals of visceral afferent fibers in the pelvic nerves and somatic afferent fibers for the pudendal nerves through the Lissauer’s tract (LT) and its lateral- and medial-collateral projections (McKenna and Nadelhaft, 1986; Thor et al., 1989; Liu et al., 1998; Bansal et al., 2017). The retrograde transganglionic labeling of primary afferent fibers from the bladder(Wang et al., 1998), urethra (Vizzard et al., 1995) and external urethral sphincter (Nadelhaft and Vera, 1996), as well as from the penile nerve(McKenna and Nadelhaft, 1986) has indicated that DGC is a part of the reflex pathways that control the functions of the pelvic viscera (Palecek and Willis, 2003; Cruz et al., 2017).

Some structure of the brain stem have a neuroanatomically reciprocal relationship with the lumbosacral spinal cord (Kuo and de Groat, 1985; Al-Chaer et al., 1996; Wang et al., 1999; Qi and Kaas, 2006; Liao et al., 2015). The DGC in the sacral segment is involved in the central processing of pelvic visceral information and is also associated with nociceptive, analgesia and autonomic function (Wang et al., 1999). With excitatory connection to the parasympathetic preganglionic neurons of the lumbosacral spinal cord, the pontine micturition center projections to the inhibitory interneurons of the sacral spinal cord, causing urination by relaxing the external urethral sphincter (Blok and Holstege, 2000; Verstegen et al., 2017). Functional evidence also indicates that the DGC receives terminations from the afferent fibers of the somatic and viscera(Al-Chaer et al., 1996).

The nicotinamide adenine dinucleotide phosphate-diaphorase (NADPH-d) reaction is used as a marker to characterize certain neuronal properties and colocalize with nitric oxide synthase (NOS) (Dawson et al., 1991; Hope et al., 1991). However, some researchers demonstrate that NADPH-d is not always identical to NOS (Belai et al., 1992; Traub et al., 1994; Pullen and Humphreys, 1995; Tan et al., 2006). In both central and peripheral nervous system, only a part of NOS positive neurons colocalize with NADPH-d histochmistry (Belai et al., 1992). Bioactivity of NOS and NADPH-d is depended on different cellular location(Matsumoto et al., 1993). Neurons with NADPH-d activity have been shown to exhibit colocalization with several neuropeptides in various brain nuclei (Spike et al., 1993; Chertok and Kotsuba, 2013). At various segmental levels of the spinal cord of the rat, NADPH-d activity is present in a large percentage of visceral afferent neurons in dorsal root ganglia (DRG) (Aimi et al., 1991; McNeill et al., 1992; Vizzard et al., 1993c; Vizzard et al., 1994b; Porseva and Shilkin, 2010). In both the rat and cat, NADPH-d is also present in a prominent afferent bundle projecting from LT to the region of the sacral parasympathetic nucleus (Vizzard et al., 1993c; a; Vizzard et al., 1994a). This afferent pathway closely resembles the central projections of afferent neurons innervating the pelvic viscera (Morgan et al., 1981; Nadelhaft and Booth, 1984; Steers et al., 1991; Vizzard et al., 1996).

Previous studies have shown that NADPH-d positive neurons and fiber networks are densely stained in the DGC of adult animals (Doone et al., 1999). In addition, studies have confirmed that an alternative neurodegenerations indicated by age-related NADPH-d positive bodies are specifically present in the lumbosacral spinal cord of aged rats (Tan et al., 2006). It could be an aging of onset and progressive marker for pelvic organ dysfunction of aging. A large number of NADPH-d positive neurons in the spinal cord appear to be involved in visceral regulation. The NADPH-d activity of the DGC and the intermediolateral column at the segments of the lumbosacral spinal cords may have a special role in the reflexes of the pelvic organs. Changes in the neurochemical properties of these neurons after a spinal cord injury may be mediated by pathological changes in the target organs (i.e., urinary, bladder) and/or spinal cord. NADPH-d positive neurons innervate most of the pelvic organs, such as the penile tissue (Tamura et al., 1994; 1995), internal anal sphincter (O’Kelly et al., 1994; Lynn et al., 1995; Chakder and Rattan, 1996), and lower urinary tract (Zhou et al., 1997; Zhou and Ling, 1998). Pelvic visceral organ-related physical and functional alterations are known to occur frequently with advancing age. The goals of the present study were to determine whether NADPH-d positive abnormality appearing in the lumbosacral spinal cord of aged dog are associated with age-related changes.

## Materials and Methods

### Animal and tissue preparation

Young (less than 2-year-old, n=8) and aged (more than 10-year-old, n=8) dogs (Canis lupus familiaris) of both sexes weighing 8–20 kg were used in our experiments. The medical records were retrieved after owners’ informed consent. These animals did not shown any neurological deficits before experiments and were humanely euthanatized. All experimental procedures were approved by the Ethics Committee in Animal and Human Experimentation of the Jinzhou Medical University. The animals were anesthetized with sodium pentobarbital (50 mg/kg i.p.) and perfused transcardially with saline followed by freshly prepared 4% paraformaldehyde in a 0.1M phosphate-buffer (PB, pH7.4). Following perfusion fixation, the spinal cords and brains were rapidly dissected out and placed in 25% sucrose for 48 hrs.

The spinal cords from the cervical to coccygeal segments and gracile nucleus in the medulla oblongata as well as the thalamus were cut transversely into one-in-three series of 40μm sections on a cryostat. To visualize the rostrocaudal orientation of the NADPH-d positivity, horizontal sections (40μm) of the spinal cords of the aged dogs were also performed. To reconfirm a part of results in the aged dogs, transverse sections of an 80μm thickness were also made.

### NADPH diaphorase histochemistry

Staining was performed using free floating sections(Tan et al., 2006). Most of the spinal cord sections from the young and old dogs were stained and examined by NADPH-d histochemistry, with incubation in 0.1 M Tris-HCl (PH 8.0), 0.3% Triton X-100 containing 1.0 mM reduced-NADPH (Sigma, St. Louis, MO, USA) and 0.2 mM nitro blue tetrazolium (NBT, Sigma), at 37℃ for 2 to 3 h. Sections were monitored every 30 min to avoid overstaining. The reaction was stopped by washing the sections with the phosphate buffered saline (PBS, 0.05M).

### Double Immunofluorescence staining

Some sections were processed by double-staining with NADPH-d histochemistry and NeuN, CGRP, VIP or GFAP immunofluorescence and single-staining with NeuN, CGRP, VIP, MAP2, Iba1 or GFAP immunofluorescence, respectively. The sections were collected in PBS in 24-well plates and processed for free-floating immunofluorescence using primary polyclonal antibodies that label neurons (NeuN, mouse; 1: 1000, Millipore MAB377, Merck Millipore), reactive astrocytes (GFAP, mouse; 1: 1000, Sigma, USA), microtubule associated protein 2 (MAP2, mouse; 1:200, Novus Biologicals, Littleton, CO, USA), calcitonin gene-related peptide (CGRP, mouse; 1:100, Sigma, USA), vasoactive intestinal peptide (VIP, rabbit, 1:1000 Sigma, USA), microglia (Iba1,rabbit; 1:1000, Wako Chemicals, Japan). Sections were incubated for 1 h at room temperature in blocking solution (0.05 M PBS) at pH 7.4 with 1% BSA. The primary antibody was diluted in PBS containing 1% BSA and applied to the sections for 24 h at 4 °C. In each immunofluorescence testing, a few of sections were incubated without primary antibody, as a negative control. The sections were then washed several times with PBS. Fluorescent-conjugated secondary antibodies (IgG anti-mouse Cy3 conjugated [1:2000, Jackson], Goat anti-Rabbit IgG (H+L), Alexa Fluor 594 [1:800, Life] and Mouse anti-Human IgG1, Alexa Fluor 488 [1:800, Life]), were diluted in PBS and applied to the sections for 1 h at 37 °C in the dark. Finally, after several washes with PBS, the sections were incubated with DAPI for 10 min. The sections were placed onto slides and coverslipped. For controls of immunofluorescence staining, the primary antibodies were omitted or replaced with the same amount of normal serum from the same species while doing the same specific labeling with normal procedure of immunofluorescence staining. No specified staining was detected the immunostaining control experiments.

### Measurement of neurons and fibers

Images were captured with a DP80 camera in an Olympus BX53 microscope (Olympus, Japan). Sections were observed under the light microscope and 20 randomly selected sections from all spinal cord levels in each animal were quantitated using Olympus image analysis software (Cellsens Standard, Olympus). The numbers of NADPH-d neurons were counted on both sides of the spinal cord, on each section of each animal. The diameter of 500 NADPH-d megaloneurites and normal fibers and the length and area of 2000 NADPH-d megaloneurites were also assistantly measured with Neurolucida 360 (MBF Bioscience, Inc, USA). Stereological analysis was also used for measurement of the NADPH-d positive cells with Stereo Investigator (MBF Bioscience, Inc, USA).

### Statistics and Figure edition

All data are expressed as the mean ± SEM and P < 0.05 were regarded as statistically significant. Statistical analyses were performed using GraphPad Prism 5.0 (GraphPad Software, La Jolla, CA). Differences between young and aged dogs of NADPH-d positive neurons in sub-regions of the cervical, thoracic, lumbar and sacral segments were analyzed using paired t-tests.

## Result

### NADPH diaphorase activity in the sacral spinal cord of aged dogs

This study primarily evaluates on the dorsal part of the sacral cord, especially the lateral collateral pathway (LCP) of LT, the sacral parasympathetic nucleus, and the DGC because of the distribution of the NADPH-d positivity. In young dogs, both fiber and cellular staining were normally detected in the dorsal horn, LT, LCP and DGC. The pattern of NADPH-d staining in the spinal cord of young dog was consistent with the previous discovery (Vizzard et al., 1997). In the aged dogs (Fig. 1A, B), a new pattern of fiber-like NADPH-d activity in the dorsal horn and DGC of the sacral spinal cord was found to be extremely different from that in young dogs (Fig. 1C, D), especially in the LCP of LT (S1-S3). An expanded-diameter fiber of non-somatic neuronal structure was detected in the sacral spinal cord in aged dog (Fig. 1E). The criteria of the aging alteration will be summarized in discussion. The swelling giant NADPH-d positive alteration was named as megaloneurite, a newly coined word, occurred in aged dogs. When compared with the young dogs, the general location of the NADPH-d positive megaloneurites and its selective segmental distribution was related to the central projection of the primary pelvic visceral sensation (black arrowheads in Fig. 1A, B), mostly located dorsal of the spinal cord. In addition, there also contained intensely-stained multipolar NADPH-d neurons with regular thinner dendrites penetrating deeply into the ipsilateral dorsal horn (open arrowheads in Fig. 1).

**Figure 1.**
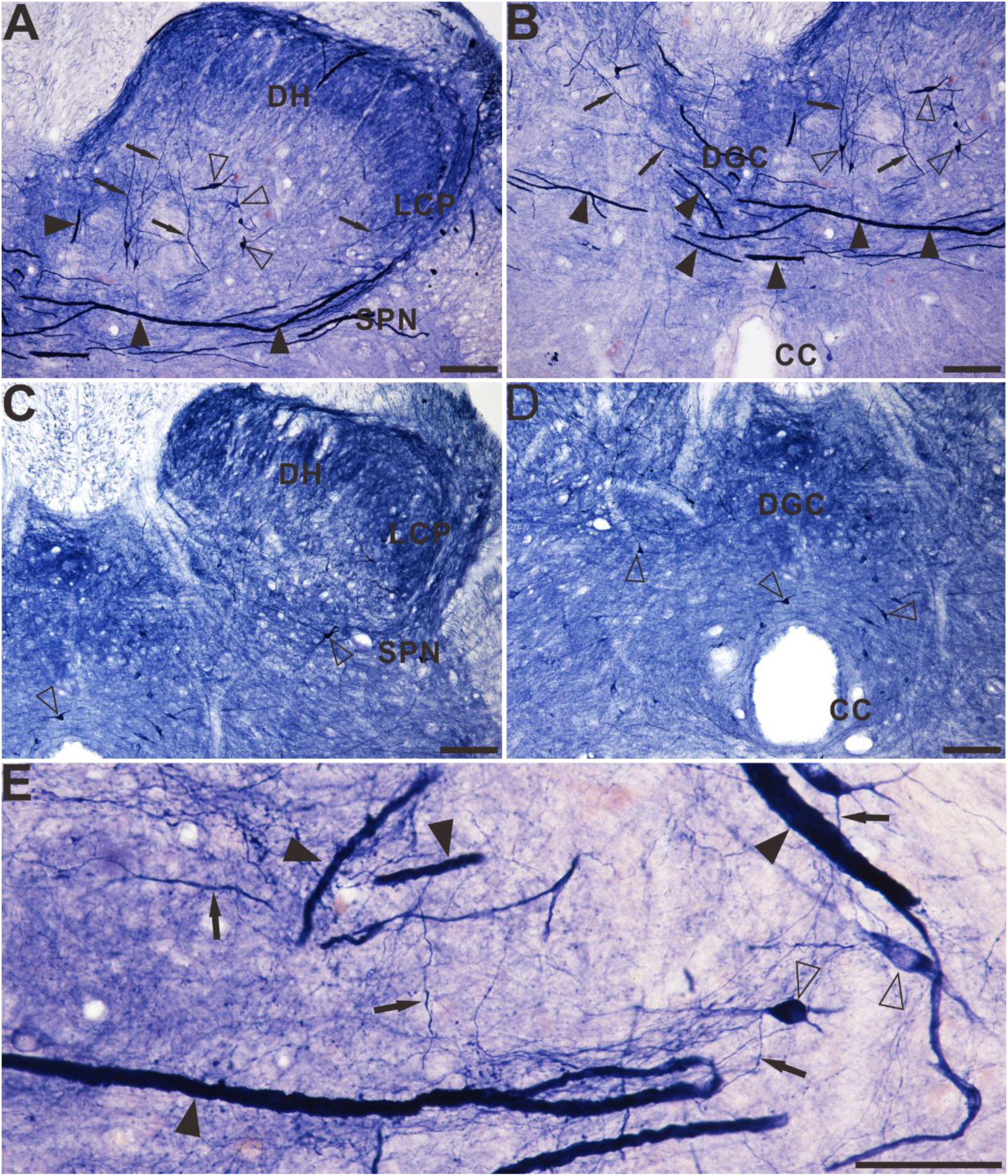
Microphotographs of NADPH-d positive reactivity in aged and young dogs at the sacral spinal cord. All of the transverse sections taken at the same levels. Note intense and abnormal NADPH-d positive megaloneurites (black arrowheads) in the LCP (**A**) and DGC (**B**) in the sacral segment of aged dogs compared with young dogs (**C** and **D**). The NADPH-d positive megaloneurites in the sacral spinal cord of aged dogs are completely different from the surrounding normal fibers and neurons (**E**). Open arrowheads: NADPH-d positive neurons, black arrows: normal NADPH-d positive neurites, black arrowheads: megaloneurites. Scale bar in **A-D**= 100μm, in **E**=50μm.

Further segmental examination was to demonstrate that the megaloneurites occurred in a specific regional localization. In transverse sections of caudal spinal segments, NADPH-d stained fiber network of dendrites and axon terminals were noted in the superficial dorsal horn (laminae I, II), the DGC, within the dorsal lateral funiculus but not in the ventral horn (Fig. 2). As mentioned above, the most prominent fiber staining in the sacral segments of aged dogs was in LT in laminae I along the lateral edge of the dorsal horn deeply into the DGC and/or passing across the DGC to the opposite gray matter (Fig. 2J). Megaloneurites were selectively detected in the sacral spinal cord (S1-S3) but not in adjacent rostral (L5-L7) or caudal (Cx1-Cx2) (Fig. 6C) segments or in thoracic and cervical segments (Fig. 2A, B).

**Figure 2.**
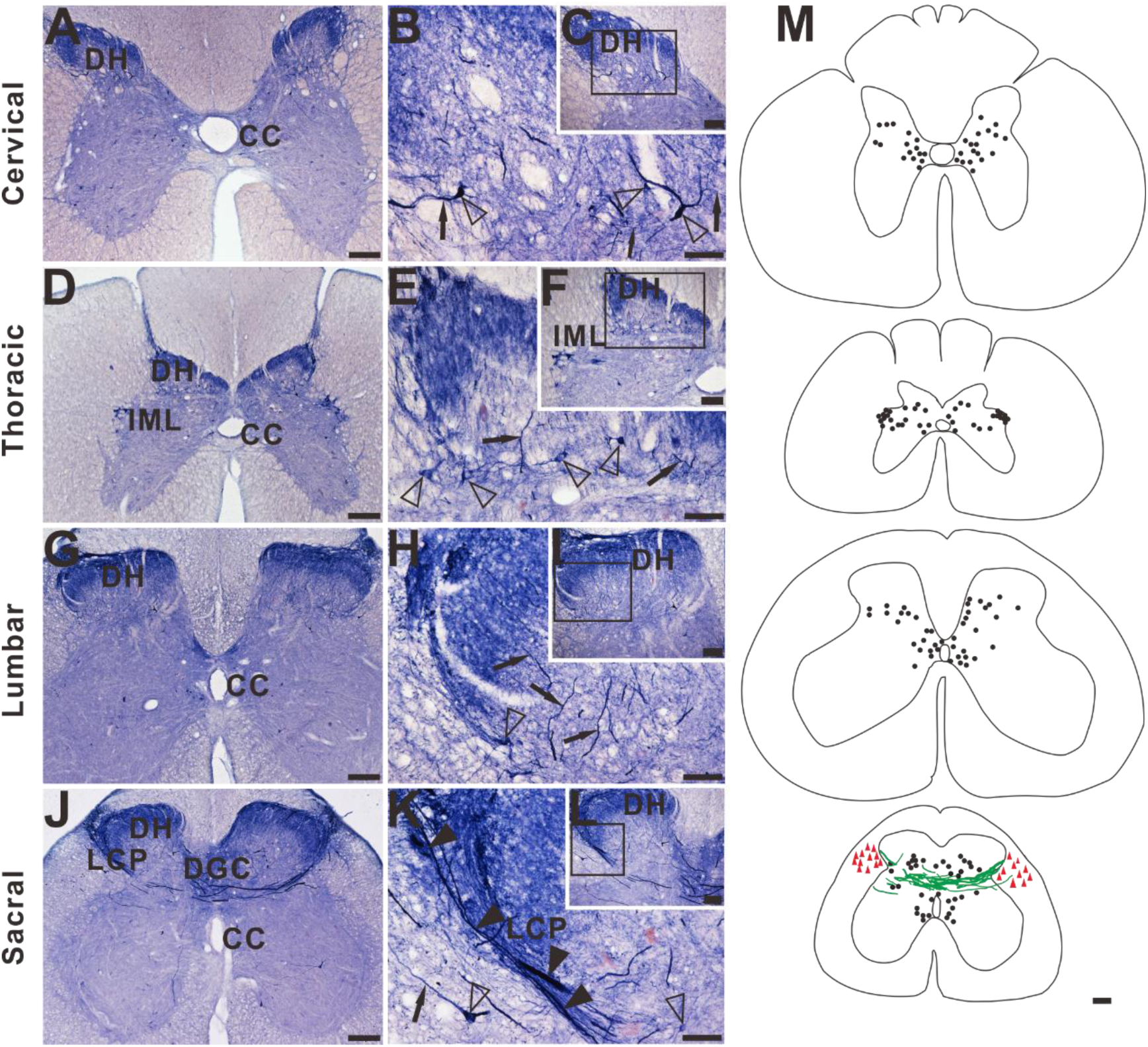
Microphotographs showing the distribution of NADPH-d staining in transverse sections of the aged dog spinal cord at different segmental levels. Few of neurons (open arrowheads) and normal fibers (black arrows) staining in the dorsal horn (DH) is present in cervical (**A**), thoracic (**D**) and lumbar (**G**) segments. Intense and abnormal fiber (black arrowheads) staining in the dorsal gray commissural (DGC) and LCP is present in sacral (**J**) segment. **B, E, H, K** show higher magnifications from insert **C, F, I, L**, respectively. (**M**) Schematic diagram of NADPH-d activity taken from the cervical, thoracic, lumbar, and sacral spinal cord segments. Neuronal cell bodies are indicated as filled circles (black) on both sides of each figure. Each filled circle represents one NADPH-d positive neuron. Dense NADPH-d stained megaloneurites are represented by cords (green). The triangle symbols (red) indicate NADPH-d activity in the white matter. NADPH-d stained neurons and fibers from 5 sections are plotted on a drawing of transverse section of the aged dog spinal cord at indicated segmental levels. Open arrowheads: NADPH-d neurons, black arrows: normal NADPH-d positive neurites, black arrowheads: megaloneurites. Scale bar in **A, D, G, J, M** =200μm; **C, F, I, L** =100μm; **B, E, H, K** =50μm.

The double-staining of NADPH-d histochemistry combined with GFAP, NeuN, CGRP, and VIP immunofluorescence were used to identify the megaloneurite properties respectively (Fig 3). No structures corresponding to NADPH-d positive megaloneurites were detected by GFAP, NeuN or CGRP immunofluorescence (Fig. 3A-I). For VIP immunoreactivity, VIP and NADPH-d mageneurites positively localized around the central canal (CC) (white arrowheads in Fig. 3J-L). In addition, VIP immunoreactivity and NADPH-d megaloneurites also had similar immunoreactions in LCP, DCN and the dorsal root entry zone (Fig.4E, F). By the immunofluorescence of NeuN (Fig. 4A, B) and MAP2 (Fig. 4C, D), it could be observed that the number of neurons in the dorsal horn of sacral segments of aged dogs was significantly decreased, and the neuronal processes were sharply reduced compared with young dogs. Similarly, the expression of CGRP and VIP in the dorsal horn and LCP of the spinal cord was also significantly reduced in aged dogs (Fig. 4G, H). These changes might be related to the degeneration of neurites in the aged. Besides, fluorescence expression of Iba1 in the sacral spinal cord of aged dogs was significantly up-regulated (Fig. 5A, B). In the superficial dorsal horn of aged dogs, the expression of fibrous astrocytes with elongated processes and less branches was sharply reduced, while the protoplasmic astrocytes with thicker processes and more branches were increased (Fig. 5C, D).

**Figure 3.**
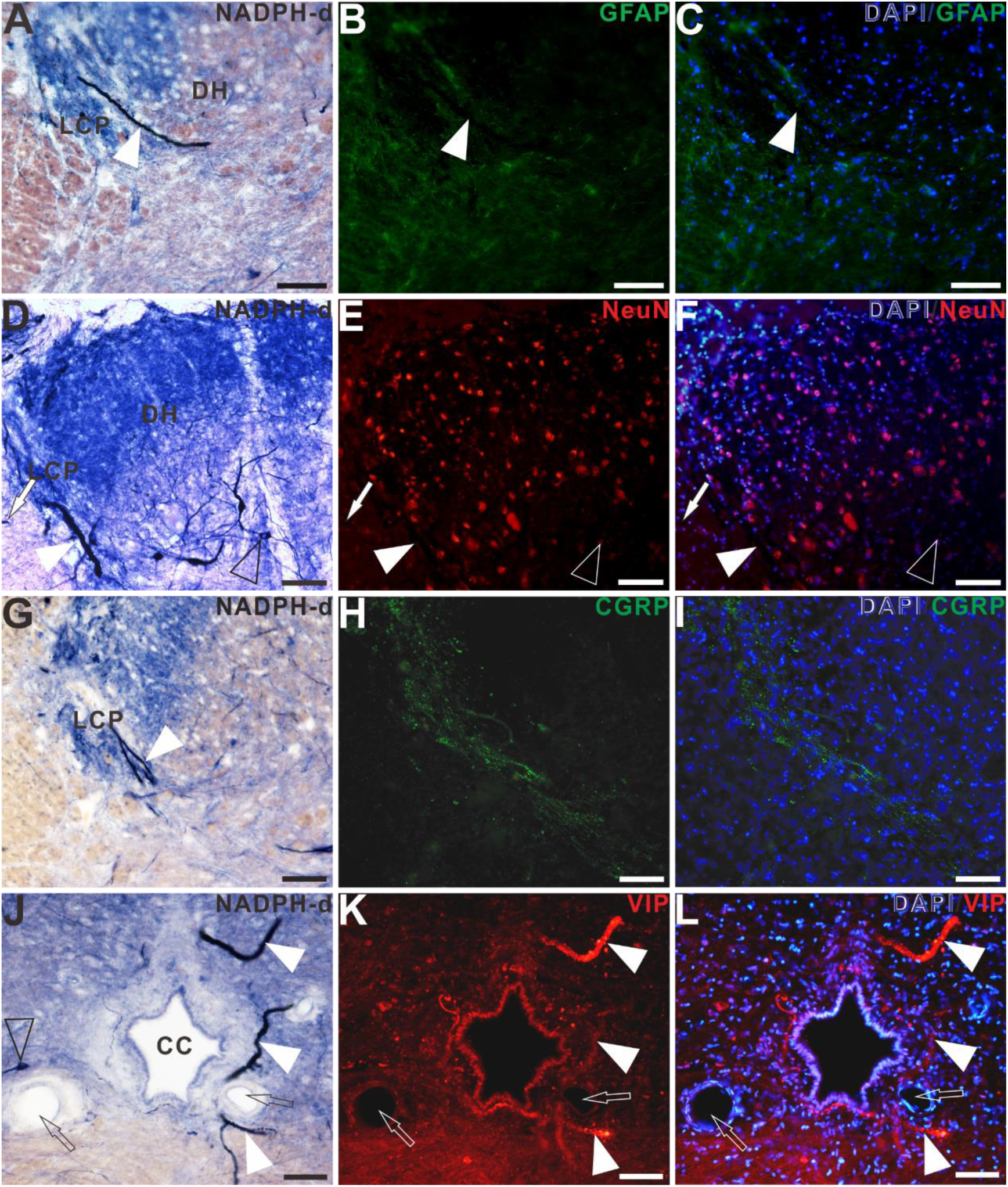
Double-staining of NADPH-d histology combined with GFAP (**A-C**), NeuN (**D-F**), CGRP (**G-I**), VIP (**J-L**) immunofluorescence in the sacral segment of aged dogs. The NADPH-d positive mageneurites (white arrowheads) in the LCP are negative for GFAP, NeuN and CGRP immunoreactivity, but positive for the VIP immunoreactivity around the CC. Open arrowheads: NADPH-d positive neurons, white arrowheads: NADPH-d positive megaloneurites, open arrows: vascular structures, white arrows: NADPH-d positive structures in the white matter. Scale bar=50μm.

**Figure 4.**
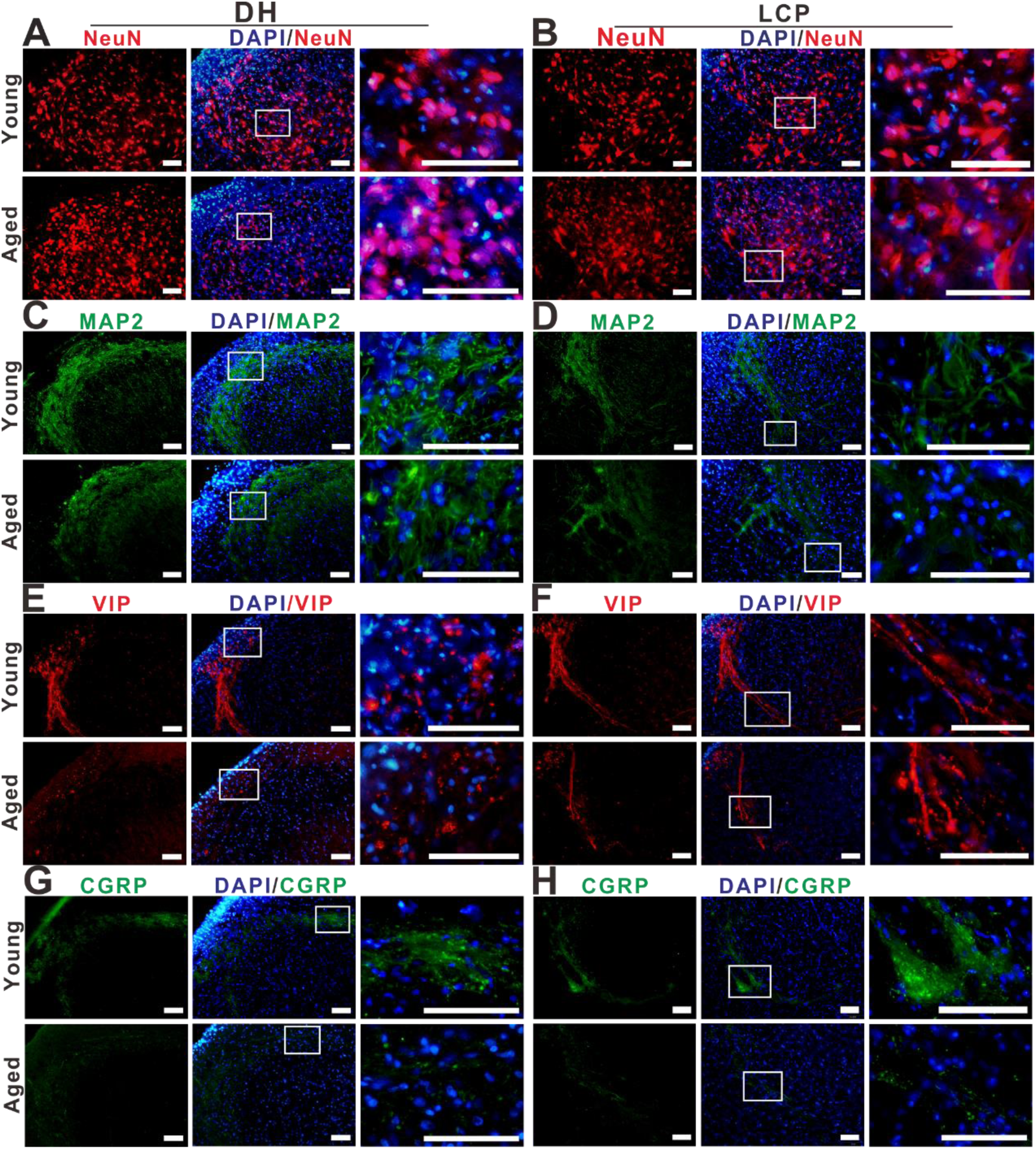
The positivity of NeuN (**A, B**), MAP2 (**C, D**), VIP (**E, F**) and CGRP (**G, H**) immunoreactivity in the DH and LCP of the sacral spinal cord of young and aged dogs. Megaloneurites indicated with VIP positive (E, P). Scale bar=50μm.

**Figure 5.**
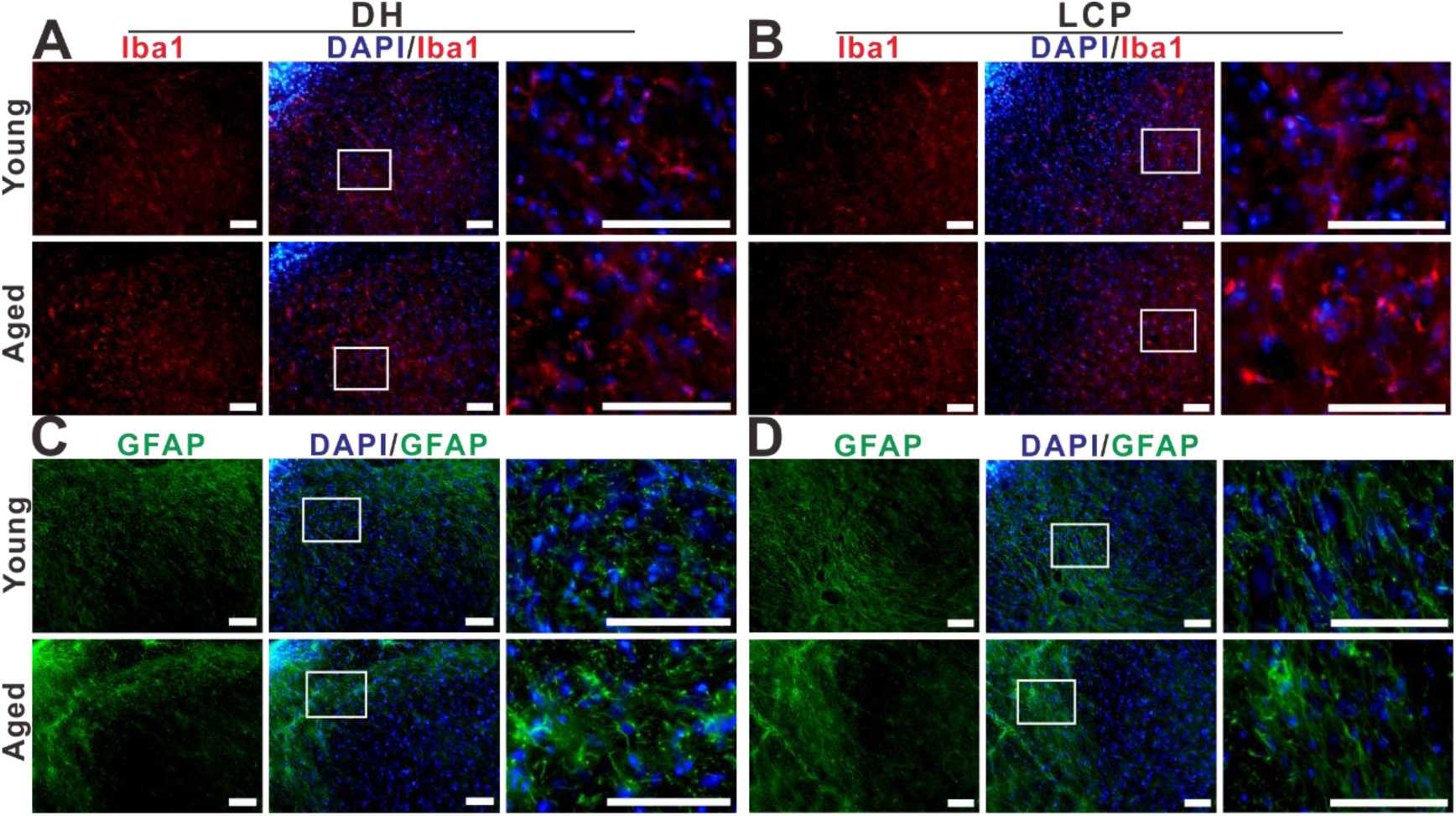
The distribution of Iba1 (**A, B**) and GFAP (**C, D**) immunoreactivity in the DH and LCP of the sacral spinal cord of young and aged dogs. Scale bar=50μm.

Statistical data indicated that the intensely stained NADPH-d megaloneurites ranging between 5 and 2296.4μm in length identified on coronal sections of thickness 40μm were regularly seen (Fig. 6A). It is noteworthy that the area of the megaloneurites ranged between 80 and 153217μm² (Fig. 6B). In the histogram, the diameter of a large proportion of the NADPH-d megaloneutrites ranged from 2 to 5μm, and normal fibers ranging from 1 to 2μm (Fig. 6C). The average diameter of the megaloneurites (4.47±0.101μm) was thicker than that of the normal fibers (1.32±0.017μm) of aged dogs and was much thicker than the NADPH-positive fibers of young dogs (1.89±0.017μm) (Fig. 6D).

**Figure 6.**
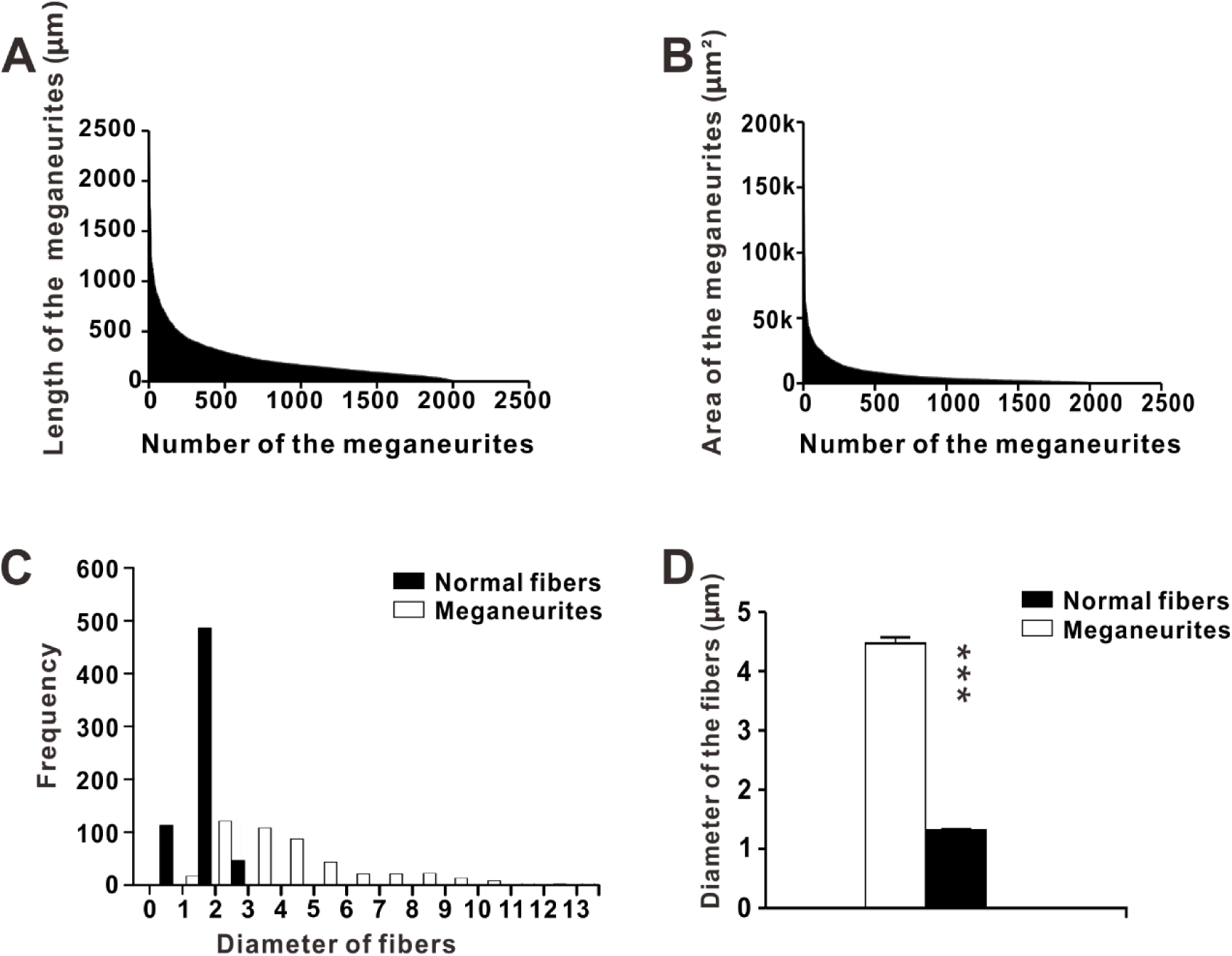
Charts showing length (**A**) and area (**B**) of the NADPH-d positive megaloneurites in the sacral spinal cord of aged dogs (2000 megaloneurites counted). (**C**) Histogram of diameter distribution of NADPH-d positive fibers. (**D**) The diameter of the megaloneurites (500 NADPH-d megaloneurites counted) and normal fibers (500 fibers counted). The k of unit label in **B** represents 1000, asterisks in horizontal bars indicate statistically significant comparisons (***P<0.0001).

The horizontal sections of the sacral cord clearly indicated the distribution pattern of the megaloneurites (Fig. 7). This spatial arrangements was correspondingly detected megaloneurites in the transverse sections in Fig. 2J, Fig1.A and B of the DGC or mediated SPN. In the horizontal sections of the sacral cord, the megaloneurites were confirmedly detected in the DGC almost vertical to along the rostrocaudal axis. The longitudinally oriented megaloneurites occurred more frequently in the LCP of LT and DGC (Fig. 7A). At higher magnifications (Fig. 7C), the size of the typical megaloneurites was much bigger than that of the regular NADPH-d positive neuronal processes (black arrows). It could be observed that the terminal of majority of the megaloneurites were branched, and the diameter of these branches (5.74±0.26μm) was significantly thinner than the diameter of the megaloneurites (12.57±0.66μm), and the diameter reduction rate reached 49%. The caliber of megaloneurites within LT and the LCP varied and included thin as well as thicker structures (Fig. 7). The interval of individual or clustered megaloneurites (Fig. 7B) was approximately 186.6±5.38μm apart, which was calculated between adjacent midpoints of megaloneurites. The pattern of arrangement was speculated that the megaloneurites in the LCP was not present in every section and it might occur intermittently along the rostralcaudal axis.

**Figure 7.**
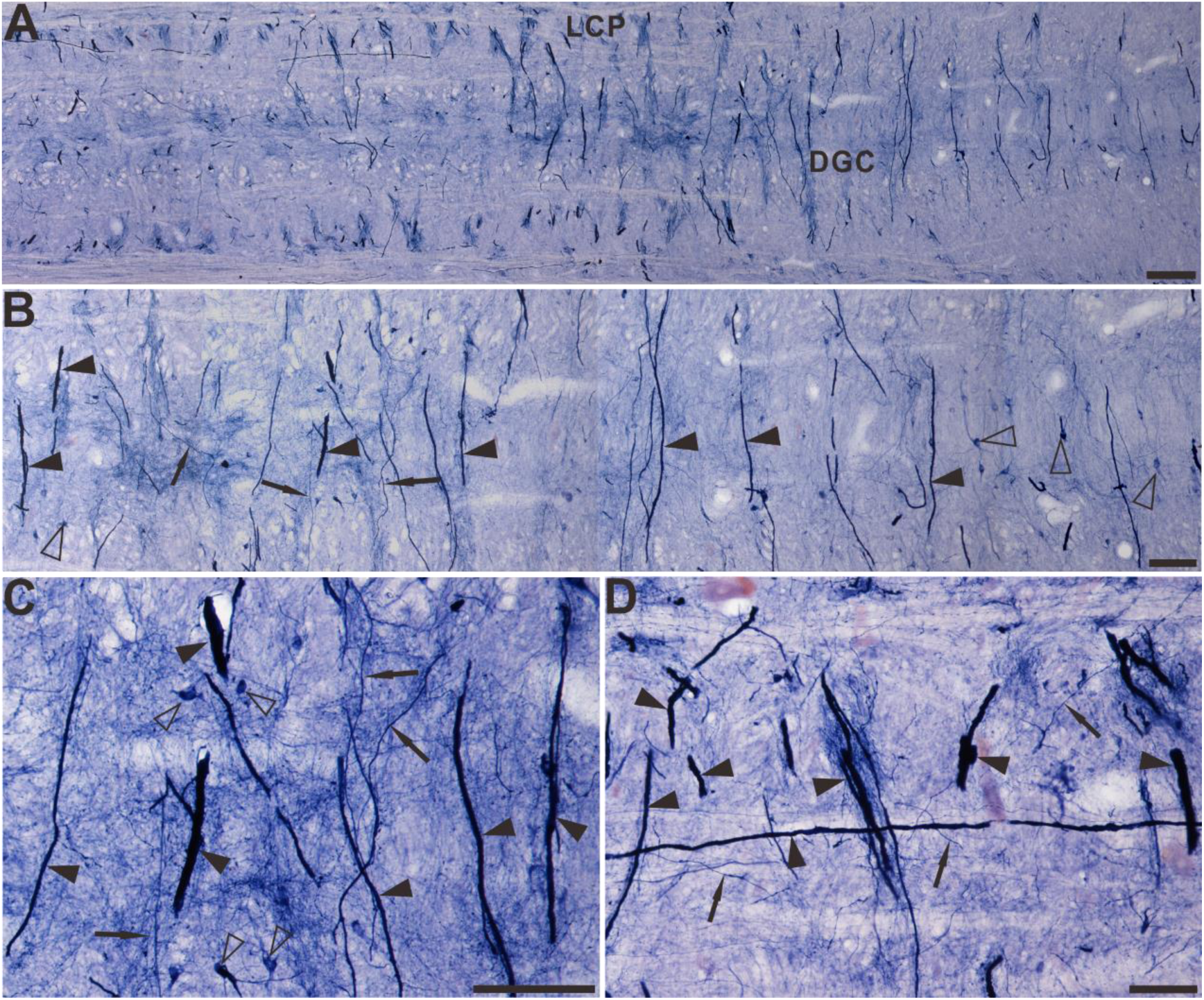
The Megaloneurites in horizontal sections in the DGC in the sacral segment of aged dogs. (A) Horizontal sections in the low power microphotograph confirmed that the megaloneurites are organized in a regular intervals vertical to the rostrocaudal axis. **B** is the magnification of **A**. (**C, D**) High power microphotograph demonstrated a significant difference between the megaloneurites and normal fibers and neurons. Open arrowheads: NADPH-d neurons, black arrows: normal NADPH-d neurites, black arrowheads: abnormal megaloneurites. Scale bar in **A**= 200μm, in **B**=100μm, in **C**, **D**=50μm.

### NADPH-d alterations around the CC in the aged sacral-coccygeal spinal cord

In contrast to the intensely-stained NADPH-d positive megaloneurites observed in the LCP at the sacral level, only a few NADPH-d positive neurons and fibers (Fig. 8A, C) were detected in the caudal sacral and coccygeal spinal segments of aged dogs. Although this megaloneurites scarcely occurred in the caudal sacral and coccygeal segments of aged dogs, one another alternative NADPH-d positive abnormality was observed around the CC of the sacral-coccygeal sections (Fig. 8). The abnormalities specifically appeared in the caudal sacral and coccygeal spinal cord of aged dogs. This NADPH-d positive abnormalities consisted of three compartments: intra-CC (Fig. 8C), inter-ependyma and extra-CC subdivisions (Fig. 8A, D). None of these profiles were found in the young dogs (Fig. 8B). It was also formed a rostrocaudal organization along with the CC in the horizontal sections. Intra-CC part attached in lumen of the CC meanwhile extra-CC one located the subcentral canal. The irregular intensely-stained NADPH-d positive abnormalities, unlike normal neurons, appeared in a mass-like or strand-like manner around the CC. The mass-like anomalous NADPH-d positive abnormalities had an area on the transverse sections ranging from 26100 to 77.15μm² and a diameter of 24.76±3.74μm. The length of these strand-like NADPH-d positive abnormalities was 117.0±8.87μm. Much enlarged NADPH-d structures could be observed in the intra-CC subdivision free of the other surrounding cellular elements (Fig. 8. C and D). In the horizontal sections (Fig. 8E-H), the NADPH-d abnormalities distributed around the subcentral canal in a strip shape and in some instances, these fiber-like structures could be traced along a whole length of the coccygeal section and could extended to 1400μm. The positive aberrant structures in intra-CC may be extracellular matrix of an amorphous profile. As mentioned above, extra-CC structures located at subependymal cellular layer as longitudinal fiber-like organizations. An anastomosis positively occurred between ependymal cells. That means intra-CC and extra-CC NADPH-d positive components were connected with an inter-ependyma subdivision through the anastomosis. We termed this NADPH-d abnormality as aging NADPH-d neuritic hypertrophy around the CC, a variation of megaloneurites.

**Figure 8.**
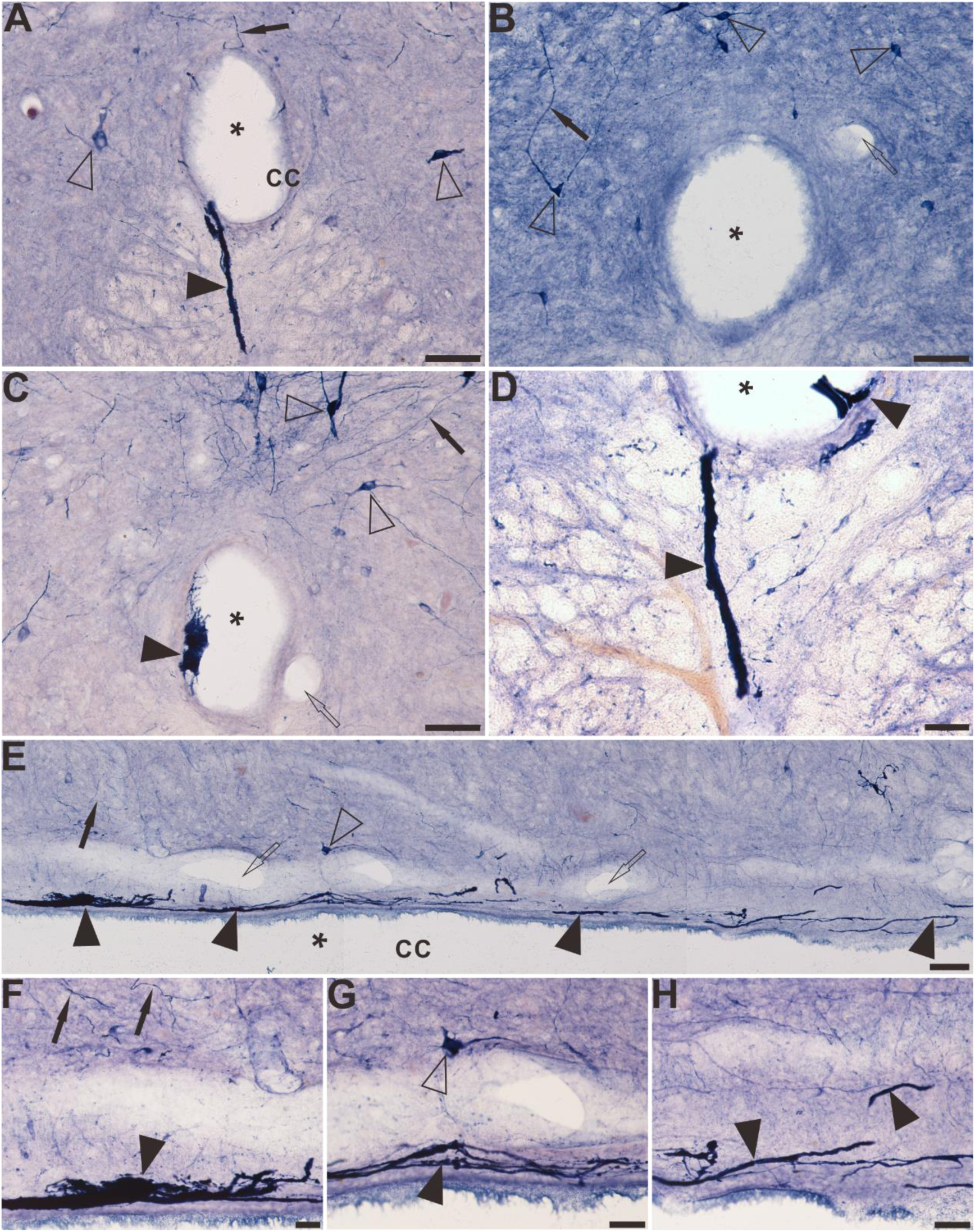
The distribution of NADPH-d positive abnormality in the sacral-coccygeal spinal cord of aged dogs. **A, C** and **D** demonstrate the location and morphology of the NADPH-d abnormalities on the transverse section of aged dogs. Some positive fibers and structures cross the ependymal cells and reach the lumen of the CC (**A**). There are no aberrant structures in the CC of young dog (**B**). Note abnormal mass-like or strand-like NADPH-d positive structures (black arrowheads) around the CC of aged dog. Both intra-CC and extra-CC alterations are detected in **D**. Megaloneurites indicated ventrally bridging from the CC to the anterior median fissure (**A** and **D). E-H** shows the location and morphology of the NADPH-d abnormalities in a remaining part of section along the CC in the horizontal section of aged dogs. Open arrowheads: NADPH-d positive neurons, black arrowheads: NADPH-d positive abnormalities, open arrows: the vascular structures, black arrows: normal NADPH-d positive neurites, the asterisk indicates lumen of CC. Scale bar in **A-C**, **E**=50μm, in **D, F-H**=20μm.

### NADPH-d activity in the white matter of aged dogs

In the sacral and coccygeal spinal cord of aged dogs, clearly-expressed punctate NADPH-d activity was noted in the white matter (see Fig.2 schematic drawing of the sacral spinal cord and Fig. 9). It was similar to megaloneurites, which may be possibly in close association with NADPH-d positive strand-like tracts penetrating deeply into the white matter (Fig. 9B). Considerably higher punctate NADPH-d activity was detected in the lateral portion of the LCP of LT, and the abnormal alterations were extremely different from normal nerve cells and neuroglia cells (Fig. 3). Statistical data indicated that the area of the punctate NADPH-d positive alterations was 43.86±1.098μm² and the diameter was 6.74±0.07μm. Horizontal sections of the sacral (Fig. 10A, C, D) and coccygeal (Fig. 10B, E) indicated that the punctate NADPH-d alterations in Fig.7 were longitudinally-arranged fibrous extending rostrocaudally for 1900-2100µm in 40-μm-thick sections and were greatly different from normal fiber bundles and vascular structures in white matter (Fig. 10). In addition, numerous NADPH-d fibers and varicosities were evident in LT and in some instances, these fibers could be traced along the whole length of the section (Fig. 10).We still termed the aged and segmental associated alterations in the white matter as megaloneurites.

**Figure 9.**
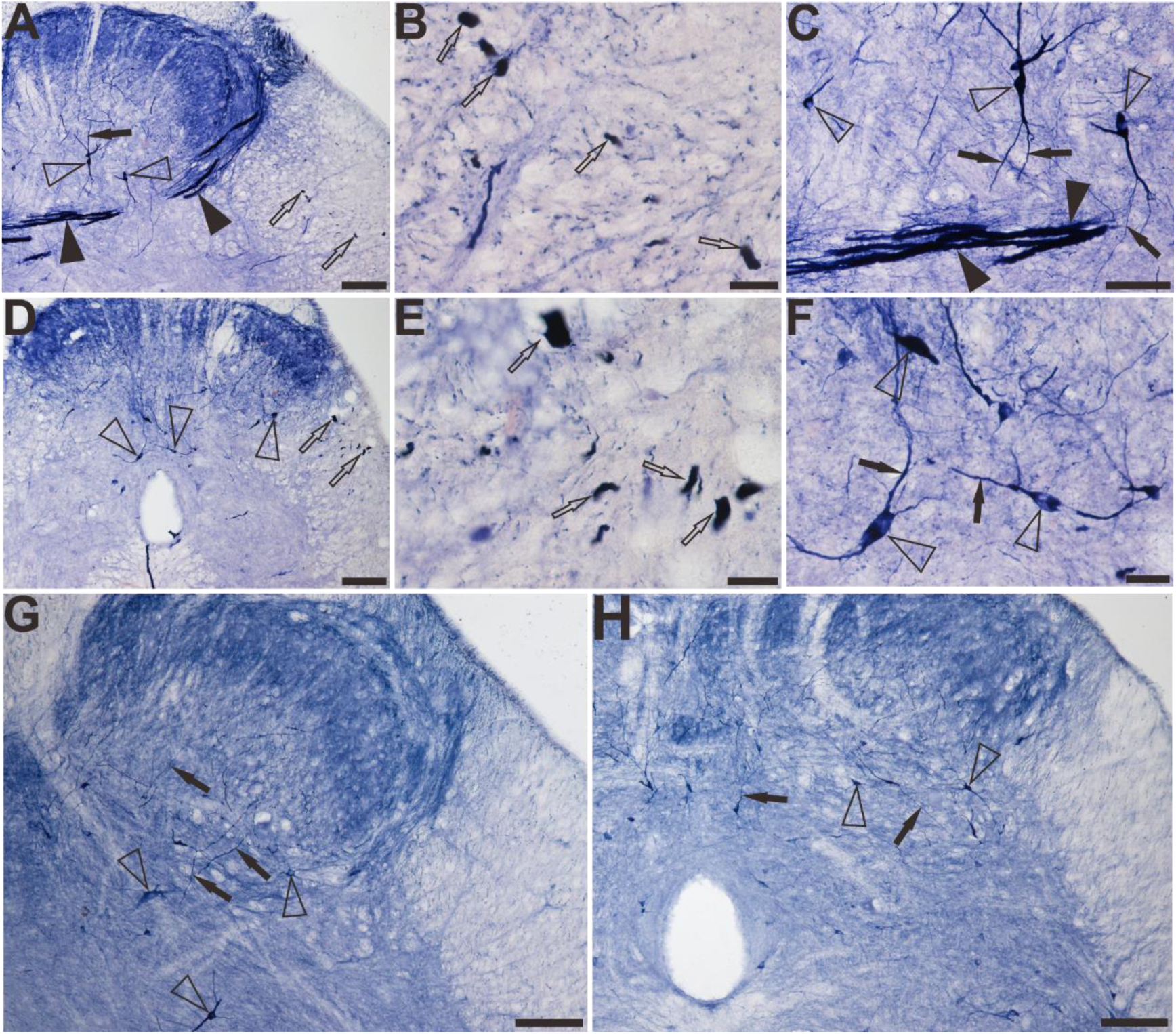
The NADPH-d positive alterations in the white matter of the sacral and coccygeal spinal cord of aged dogs. **A-C** indicate that the location and morphology of the megaloneurites in the white matter in sacral segment of aged dogs and (**D-F**) represent the coccygeal segment. **G** and **H** represent the sacral and coccygeal spinal cord of the young dogs respectively. Open arrowheads: NADPH-d positive neurons, open arrows: the alterations of NADPH-d activities detected as transection of megaloneurites (see Fig.10) in the white matter. Black arrowheads: the NADPH-d megaloneurites, black arrows: the normal NADPH-d positive neuritis. Scale bar in **A, D, G, H**=100μm, in **C, F**=50μm, in **B**, **E**=20μm.

**Figure 10.**
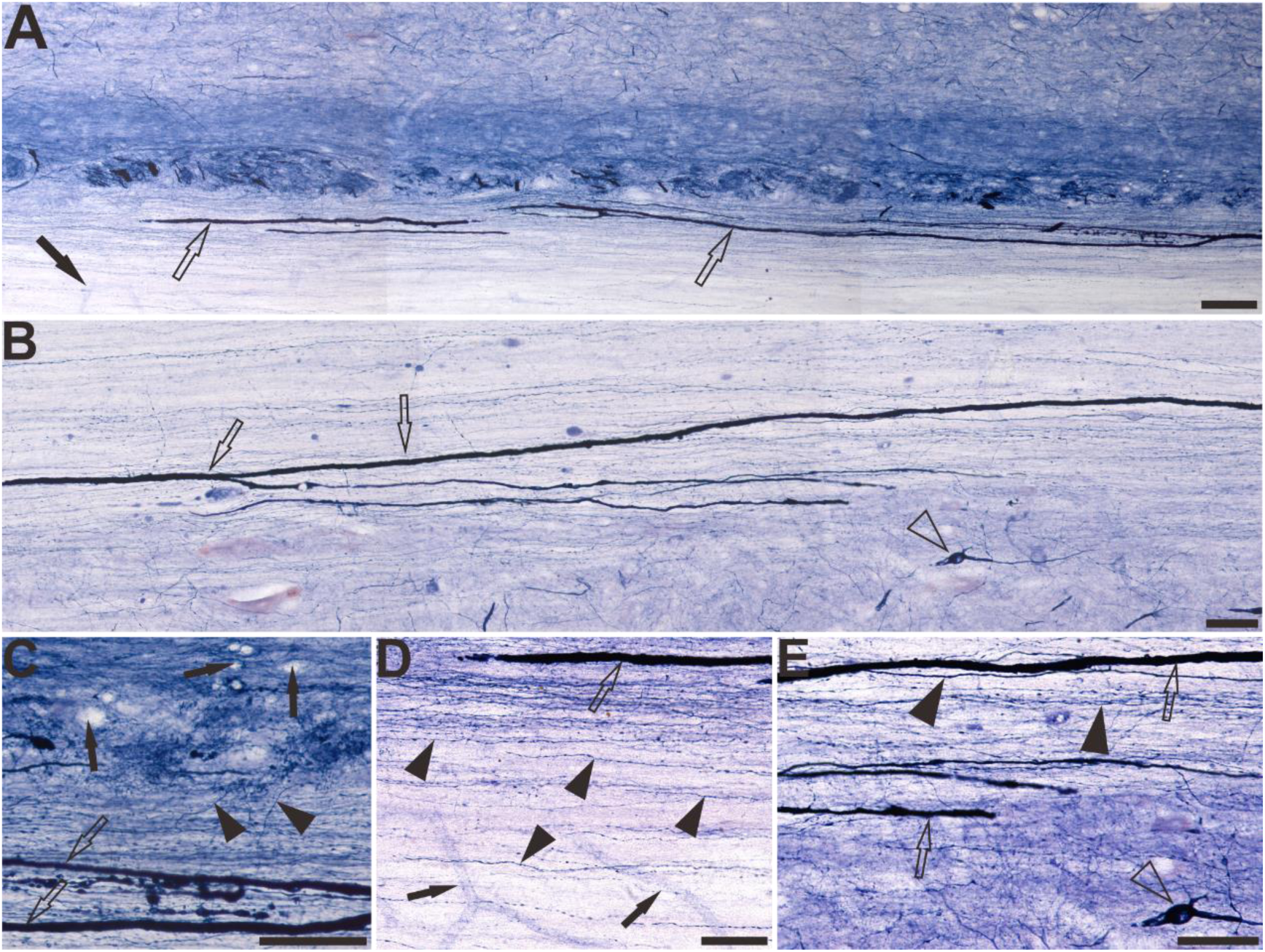
In the horizontal section, the white matter of the lateral fasciculus of the aged dogs. The NADPH-d positive alterations in the white matter of the sacral **(A, C, D**) and coccygeal (**B, E**) spinal cord. Open arrowheads: NADPH-d positive neurons, open arrows: the megaloneurites in the white matter, black arrowheads: normal NADPH-d positive neurites, black arrows: vascular structures. Scale bar in **A, B**=100μm, in **C, D, E**=50μm.

### NADPH-d activity in the caudal medulla

In the caudal medulla, primarily in the gracile nucleus and cuneate nucleus (Fig. 11), no NADPH-d positive abnormality appeared. Small, moderately stained NADPH-d positive neurons were detected both in the gracile nucleus and cuneate nucleus of the caudal medulla of the aged and young dogs. In the double-stained sections of NADPH-d histochemistry combined with GFAP and NeuN immunofluorescence (Fig. 12), there were three sub-grouped neurons: both single labeling of the NADPH-d positive neurons (black arrows in Fig. 12) and NeuN immunofluorescent neurons (white arrowheads in Fig. 12), and double-labeling neurons (open arrowheads in Fig. 12). The second order sensory axons project to the thalamus. We also examined the NADPH-d activity of the ventral posterolateral nucleus of the thalamus receiving the second-order sensory ascending projection from the dorsal column nuclei, and no positive abnormal alterations occurred (data not shown here).

**Figure 11.**
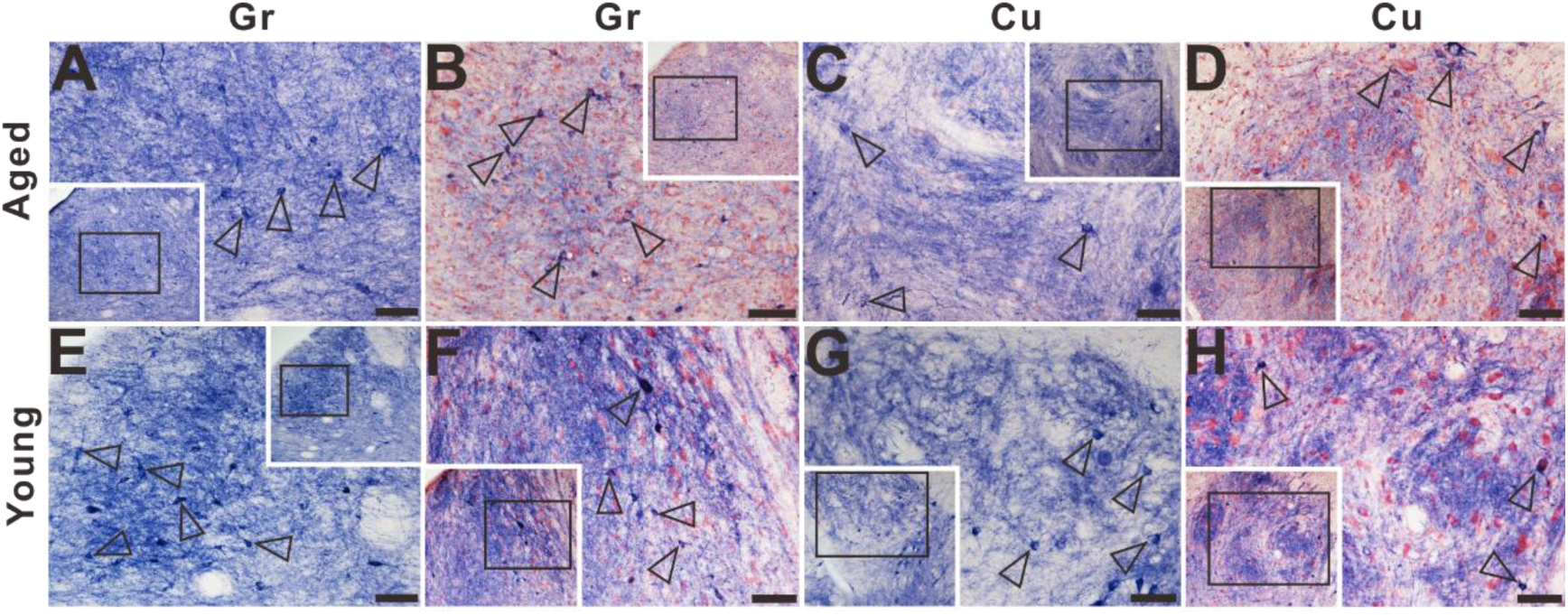
Microphotographs of NADPH-d positive reactivity in the dorsal column nuclei cournterstained with neutral red in aged (**A-D**) and young (**E-H**) dog in the gracile nucleus and cuneate nucleus. **A, C, E, G** show higher magnifications from corresponding inset respectively. No typical NADPH-d positive alterations were detected in gracile nucleus and cuneate nucleus of aged dogs compared with young dogs. Open arrowheads: NADPH-d positive neurons. Scale bar =50μm

**Figure 12.**
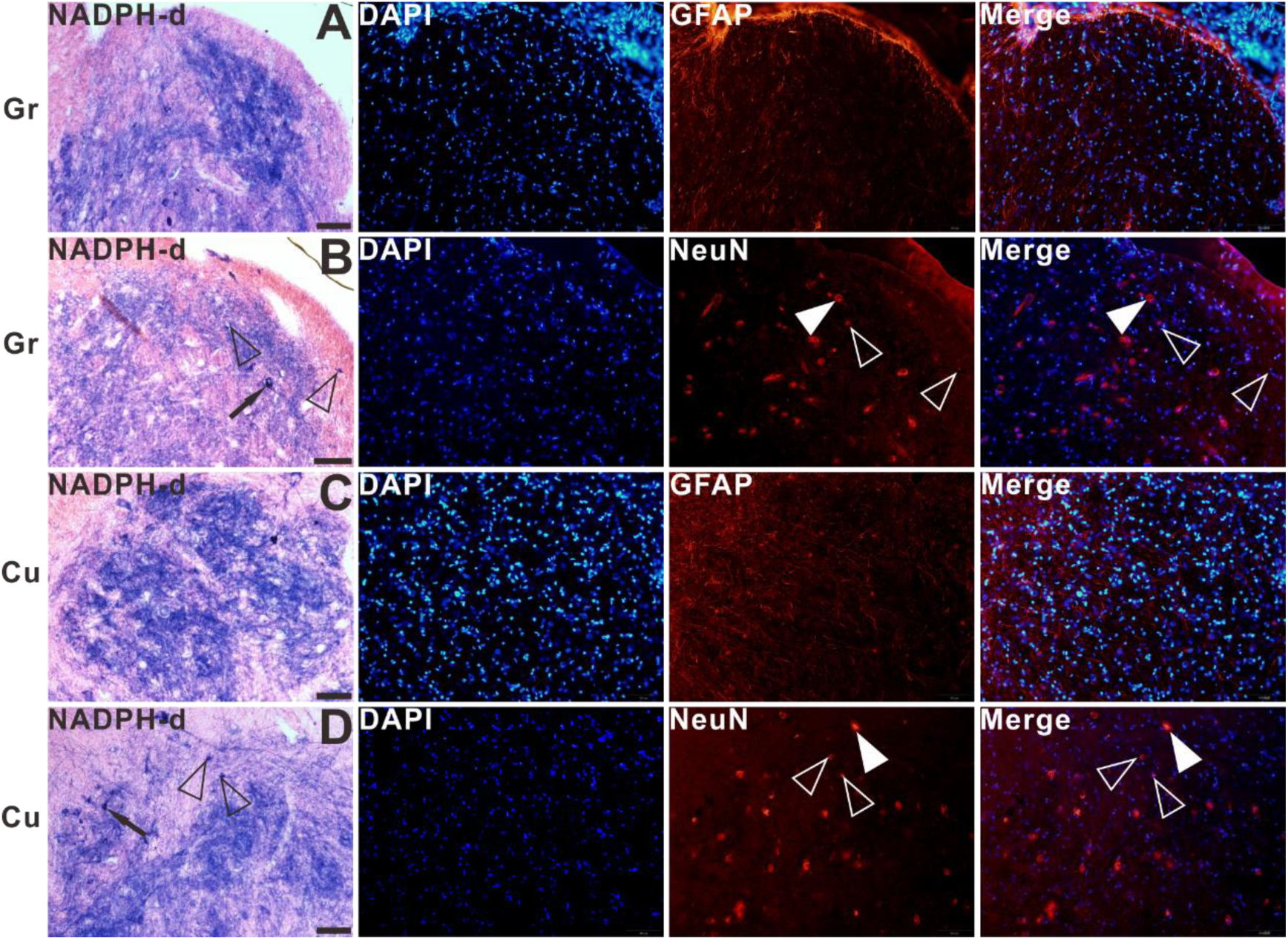
The double-staining of NADPH-d histochemistry combined with GFAP or NeuN immunofluorescence in gracile nucleus and cuneate nucleus of aged dog. Open arrowheads: double-labeled neurons, white arrowheads: single labeled NeuN immunofluorescent neurons, black arrows: single labeled NADPH-d positive neurons. Scale bar=50μm.

### Changes of NADPH-d positive neurons in the spinal cord at different segments

The changes of NADPH-d positive neurons at different segments in young and aged dogs were examined (Fig. 13). Our data showed that the number of NADPH-d positive neurons was significant reduced (P<0.05) in the dorsal horn (DH), ventral horn (Gravholt et al.), around the CC at all levels of aged dogs compared with young dogs. Such a decrease, however, was not observed in the IML of the thoracic segment of aged dogs, where the number of NADPH-d neurons did not change (P=0.94) compared with young dogs. The number of NADPH-d neurons in IML in aged dogs (15.4±1.07 cell profiles/section) were similar to those seen at the young dogs (15.5±0.84 cell profiles/section). Accompanied by aging, the NADPH-d neurons in the lumbosacral spinal cord of the aged dogs we studied had a significant decrease (P<0.01). In addition, a group of large, lightly stained motoneurons (Fig. 14) were observed in the ventral horn. These neurons were different from intensely stained neurons. It was clear from the statistics that the number of motoneurons in the ventral horn of aged dogs was significantly increased in the thoracic and lumbar spinal cord compared with young dogs (P<0.0001), but there was no significant difference in the cervical and sacral spinal cords.

**Figure 13.**
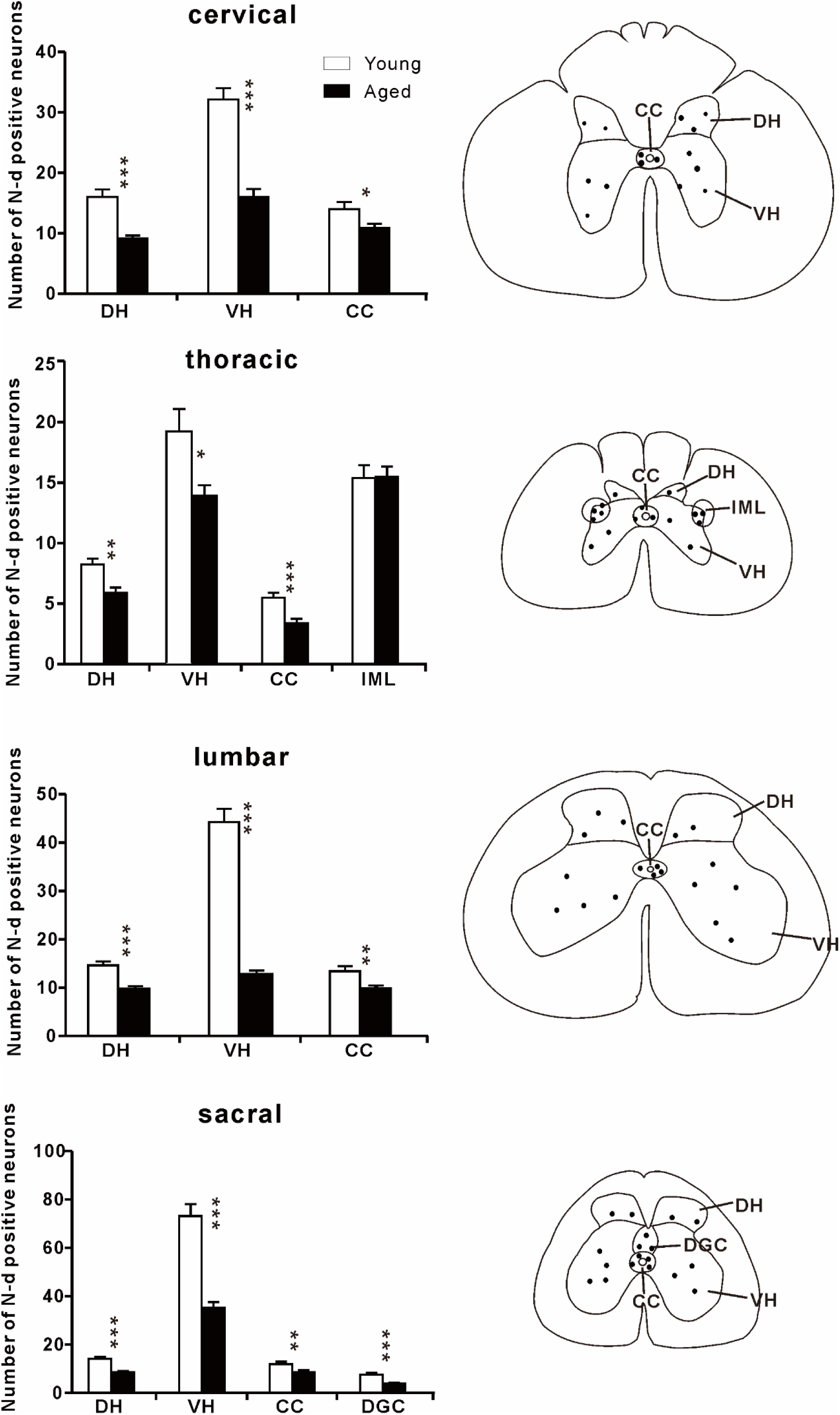
Aging changes of the NADPH-d neurons in the spinal cord. Left panel: histograms showing the number of NADPH-d positive neurons per 40-μm thickness section in sub-regions of the cervical, thoracic, lumbar and segments of the dog’s spinal cord (20 sections examined). Right panel: for specialization areas used in left panel, representative schematic diagrams based on neurolucida drawings showing the locations of NADPH-d positive neurons (black dots) derived from corresponding sections of the spinal cord. DH: dorsal horn, VH: ventral horn, CC: central canal, IML: intermediolateral cell column, DGC: dorsal gray commissure. Asterisks in horizontal bars indicate statistically significant comparisons (*p < 0.01; **p < 0.001; ***p < 0.0001).

**Figure 14.**
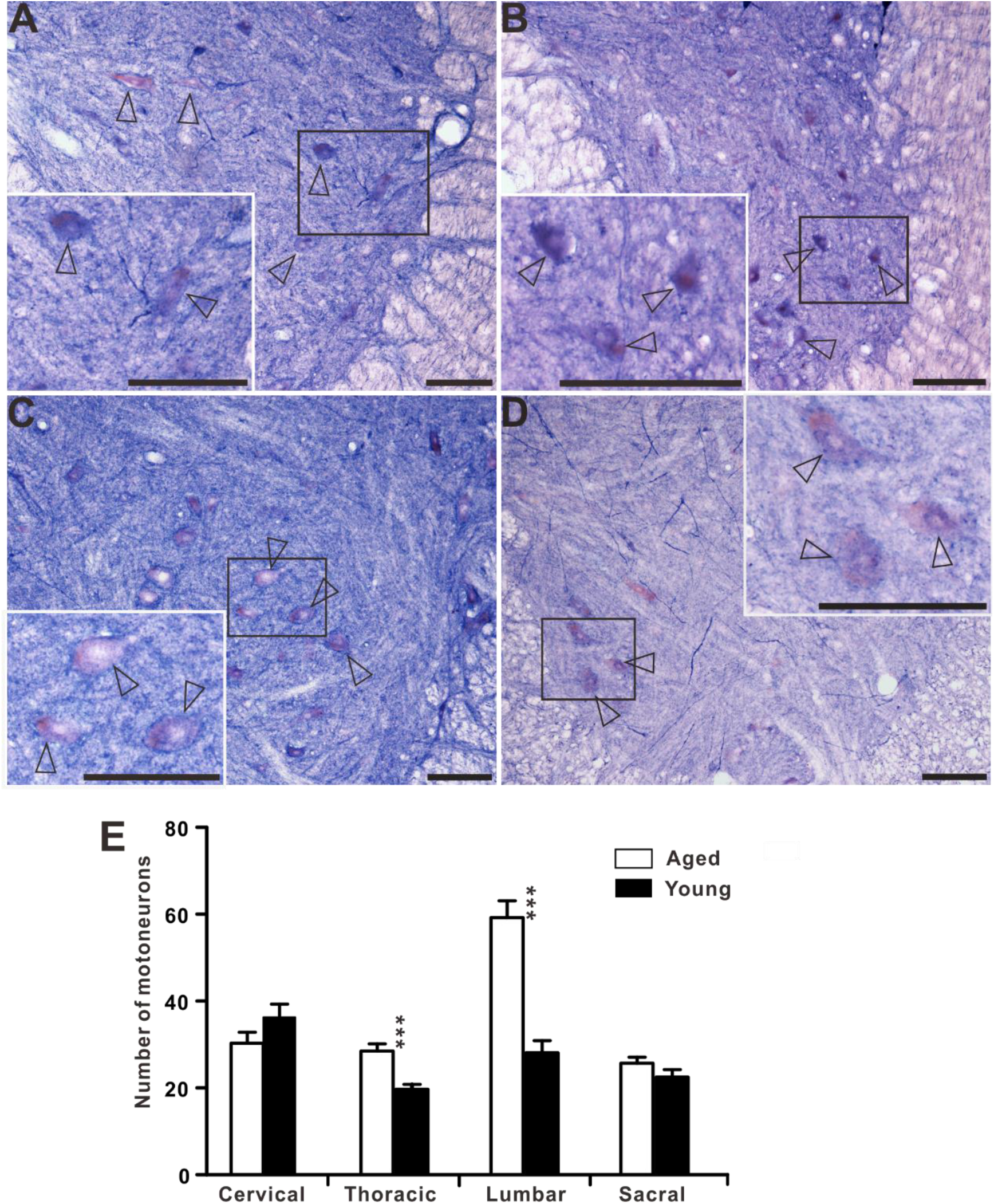
The large, lightly stained motoneurons in the transverse sections in the ventral horn of cervical (**A**), thoracic (**B**), lumbar (**C**) and sacral (**D**) spinal cord of aged dogs. (**E**) Histograms showing the number of motoneurons per 40-μm thickness section in sub-regions of the cervical, thoracic, lumbar and segments of the dog’s spinal cord (20 sections examined). Open arrowheads: large, lightly stained motoneurons. Asterisks in horizontal bars indicate statistically significant comparisons (***p < 0.0001). Scale bar=100μm.

### NADPH-d activity in the DRG and dorsal root entry zone

In the sacral DRG, cells and fibers exhibited NADPH-d activity (Fig. 15A, B). The small DRG cells exhibited the most intense NADPH-d activity, whereas the large cells were unstained or lightly stained. These results similarly coincide with those obtained from previous studies of the DRG (Vizzard et al., 1994a; Vizzard et al., 1994c). In addition, numerous NADPH-d axons, some of which exhibited varicosities, occurred throughout the ganglia. Transverse section in the dorsal root entry zone at sacral segment, showing numerous dot-like intensely NADPH-d activities accompanied with the NADPH-d megaloneurites in the LCP. In horizontal sections, NADPH-d reaction product was identified in the incoming rootlets and within the cord in the dorsal root entry zone associated with the fine fibers which are continuous with LT (Fig. 15E).

**Figure 15.**
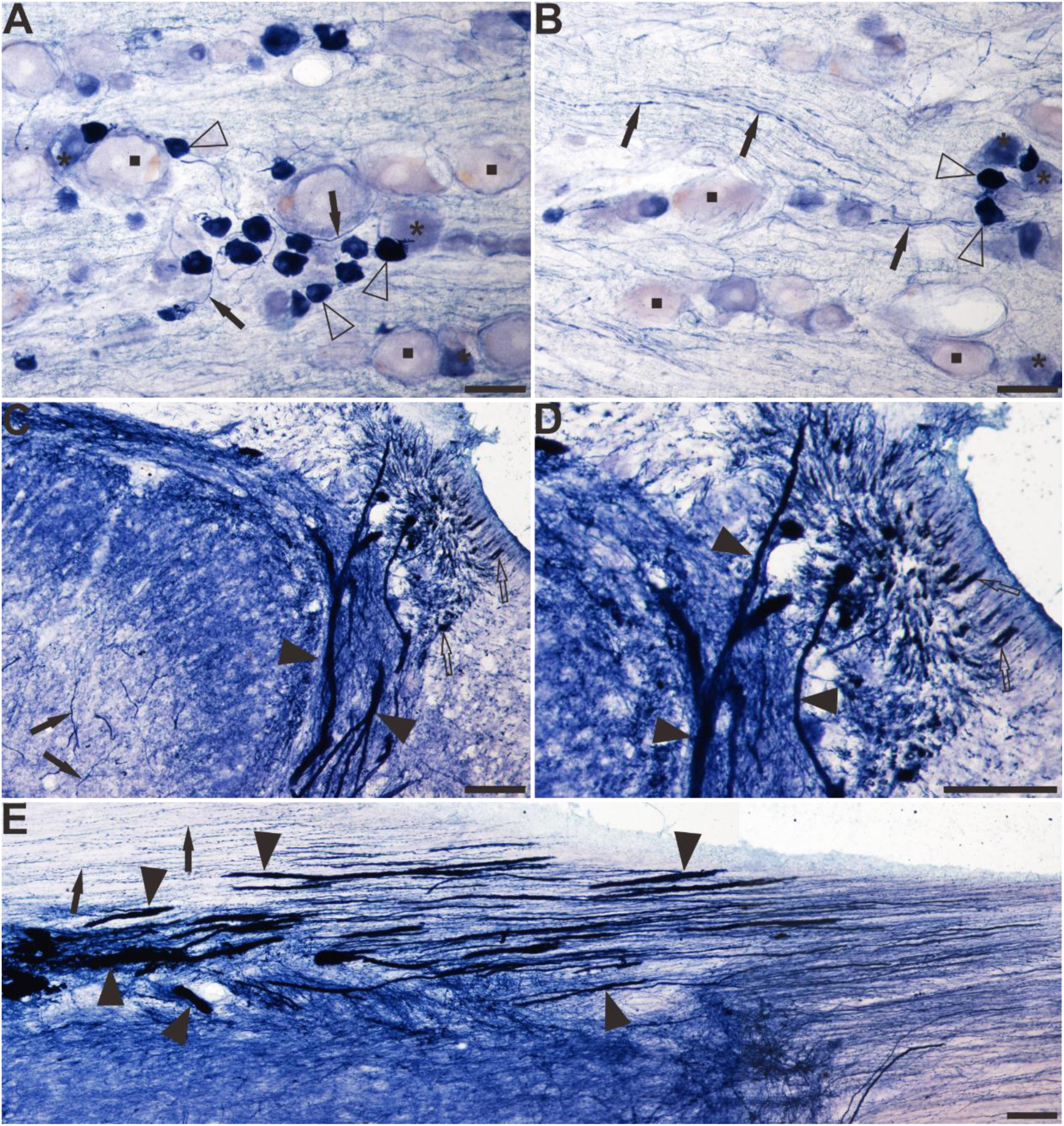
The distribution of NADPH-d positive somata and fibers in the DRG and dorsal root entry zoon of the sacral spinal cord of aged dogs. (**A, B**) A longitudinal section through sacral DRG. Intensely stained NADPH-d-positive cells (open arrows) were observed in each ganglia. (**C, D**) Transverse section in the lateral edge of the dorsal horn at sacral segment, showing LT filled with NADPH-d activity (open arrows) from its lateral border to beneath the dorsal root (DR). (**E**) Horizontal section at the sacral segment level showing NADPH-d positive neurites and aberrant megaloneurites in the dorsal root entry zone, transitionally continuous with LT. Open arrowheads: intensely stained NADPH-d positive cells, black arrowheads: megaloneurites, open arrows: dots-like NADPH-d activities, black arrows: normal NADPH-d positive neurites, the asterisk indicates lightly stained moderate-size cells, the quadrate indicates unstained large cells. Scale bar in **A, B, D**, **E**=50μm, in **C**=100μm.

## Discussion

Similar to Golgi-impregnated staining, NADPH-d histology can visualize the neuronal processes. NADPH-d reactivity occurs extensively in the spinal cord neurons and sensory pathway (Anderson, 1992; Valtschanoff et al., 1992; Vizzard et al., 1993b; Vizzard et al., 1994a; Vizzard et al., 1997; Marsala et al., 1998; Doone et al., 1999; Kluchova et al., 1999; Marsala et al., 1999). Our previous study shows that an age-related NADPH-d neurodegeneration occurred in the lumbosacral spinal cord of aging rats(Tan et al., 2006). Although the normal and some experimental NADPH-d positivity are studied in the dog spinal cord, little report is available on the aging related changes (Vizzard et al., 1997; Marsala et al., 1998; Orendacova et al., 2000).

The major of this investigation is an aged-related NADPH-d positive alterations we term as megaloneurites in the lumbosacral spinal cord of the aged dogs, and their basic morphology, segmental and laminar distribution were examined. We created the new word “megaloneurite” specialized referred megacolon and megamitochandria. The profile of the megaloneurites were postulated as dilation or swelling neuronal processes or neurites and a kind of neurodegenerative structures. This aging alterations of neuronal processes are related as a hypertrophy condition. Regular NADPH-d positive fibers stained with clear puncta and numerous varicosities. Megaloneurites are strongly stained with less punctate and varicosity, but clearly traceable for considerable distances in regular existing neuroanatomy structures especially confirmed in the horizontal sections. Staining of blood vessels is relatively weak for endothelium. The distribution of blood vessels individually separates and hardly forms bundles and tracts. We classification of the megaloneurites or abnormality into four major subcategories according to their anatomical positions: (1) megaloneurites in gray matter, (2) megaloneurites in white matter, (3) megaloneurites in dorsal entry zone, and (4) aging NADPH-d neuritic hypertrophy around the CC.

While the histologically observed segmental distribution of NADPH-d activity in the dog is comparable to that described in the spinal cord of other species such as rat (Aimi et al., 1991; Anderson, 1992; Valtschanoff et al., 1992; Saito et al., 1994), and cat (Vizzard et al., 1994a; Vizzard et al., 1994c), striking differences were noted in the LCP of LT and DGC mainly in the lower lumbar and sacral segments. In addition, the sympathetic autonomic nucleus (IML) in the rostral lumbar segments exhibited prominent NADPH-d cellular staining whereas the parasympathetic nucleus (SPN) in the sacral segments was not well stained (Vizzard et al., 1994a). The NADPH-d positive megaloneurites with morphology and organization were first discovered in the sacral spinal cord in aged dogs. Our study revealed a special occurrence of the aged-related NADPH-d positive megaloneurites in dorsal part of the lumbosacral spinal cord. The dense hypertrophic megaloneurites extending from dorsal entry zone and LT through lamina along the lateral edge of the dorsal horn to the region of the sacral parasympathetic nucleus (SPN). The sacral DGC and the LCP receives terminations from the somatic and visceral afferents (Morgan et al., 1981; Thor et al., 1989; Al-Chaer et al., 1996; Bansal et al., 2017). Functionally, the sacral spinal cord is known to be associated with bowel bladder, and sexual dysfunction (Cohen et al., 1991; Ogiwara and Morota, 2014; Barbe et al., 2018). Previous studies have demonstrated (Santer et al., 2002) that the sympathetic preganglionic neuron populations that project into the major pelvic ganglion, and the spinal inputs that they receive, exhibit a number of degenerative changes in aged rats (24 months old) which were not seen in the parasympathetic preganglionic neuronal populations. However, the distribution of the megaloneurites overlapped both the efferent and afferent pathways of the autonomic system, which regulates the pelvic organs.

In our experiments, that the megaloneurites were most prominent in sacral spinal segments (S1-S3) extending from LT to the region of the sacral parasympathetic nucleus and the DGC. These fibers may play a role in visceral reflex pathways (Morgan et al., 1981; Nadelhaft and Booth, 1984). Dorsal-ventral rhizotomy eliminated fiber staining in LT and the LCP in both the cat and the rat, but did not alter staining in the dorsal commissure or dorsolateral funiculus (Vizzard et al., 1993c). This indicated that fibers staining in LT and the LCP reflect afferent projections. Certain aspects of the NADPH-d staining are similar to that described in the rat spinal cord, although there are also differences in staining between the two species. One major difference between NADPH-d staining in rats and dogs was noted in fiber in the LT and the LCP on the lateral margin of the dorsal horn in the sacral spinal cord. In the dogs, a prominent group of NADPH-d positive fibers travelled rostrocaudally in LT and sent projections dorsoventrally along the lateral edge of the dorsal horn into the base of the dorsal horn where they turned medially. In contrast, the abnormal NADPH-d fiber bundles or megaloneurtites were not revealed in the aged rats but rather appeared spherical aged-related NADPH-d positive bodies in the sacral DGC and dorsal horn (Tan et al., 2006). The other prominent difference in NADPH-d activity between dog and rat was the lack of staining of cells in the region of the SPN of the dogs. In the rats a large percentage of preganglionic neurons in the region of the SPN was stained, while a few scattered cells in the dogs were NADPH-d positive. No enlarged fibers were found in the aged rat white matter. No aberrant NADPH-d changes were found around the aged rat CC either.

The megaloneurites were confirmedly detected in horizontal sections as a discontinuous fiber bands extending mostly in the transverse plane, and the fibers exhibited a periodicity with an interval of approximately 180μm. On the basis of these profiles, we suggest that they are a part of the neurites in the lumbosacral dorsal spinal cord. As for their origin, the question arises as to whether any other components of the age-related structures came from the glia or blood vessels. The NeuN immunofluorescence failed to label the megaloneurites. On one hand, NeuN immunoreactivity could be detected in nuclei, perikarya, and some proximal neuronal processes (Wolf et al., 1996), on the other hand, it weakly revealed in the spinal cord of aged rats (Portiansky et al., 2006). The aging-related megaloneurites in aged dogs were clearly distinguished from the glia and endothelium, as the size of typical ones was evidently bigger than the regular glia and they are also significantly different from the vascular structures found under the light microscopy(Tan et al., 2006). Testing with other serial antibodies, double staining test indicated that the megaloneurites were also VIP positive structures.

Like NADPH-d fiber staining, pelvic visceral afferent projections also exhibit a periodic distribution in the LCP of the sacral dorsal horn and extend medially from the LCP in small bundles through laminae V, VI, and VII toward the CC (Vizzard et al., 1994a). Previous tracing experiments in the rat have also demonstrated that a large percentage of afferent neurons projecting to pelvic visceral organs (McNeill et al., 1992; Vizzard et al., 1993b) and specifically to the bladder and urethra (McNeill et al., 1992; Vizzard et al., 1993c) exhibit NADPH-d activity. It has been speculated that the megaloneurites may be related to the degeneration of visceral afferent fibers in the aging condition. Horizontal spinal sections of aged dogs also provided clear evidence that megaloneurites occurred in the lateral white matter in the caudal sacral spinal cord. In the adult rabbit spinal cord white matter, sizes of NADPH-d reactive axons vary widely. Maximal NADPH-d axons at 2.5-3.5μm in diameter are identified in the sulcomarginal fasciculus as a part of the ventral column in the cervical and upper thoracic segments and in the long propriospinal bundle of the ventral column in Th3-L3 segments (Marsala et al., 2003). In the same experiment, the thinner NADPH-d axons (0.3-0.5μm in diameter) in the white matter is detected in large number within LT, while another group of NADPH-d axons (1.0-1.5μm in diameter) occurred in the LT. We found that the average diameter of megaloneurites is three times over that of regular NADPH-d fibers in the aged dogs (megaloneurites 4.47±0.101μm vs. normal fibers 1.32±0.017μm).

The megaloneurites evidently occurred dorsal root entry zone by horizontal sections of aged dogs. Analysis of the horizontal sections, diameter of neurites are gradually enlarged from distal to proximal parts through the dorsal root entry zone. The interface between peripheral nervous system and CNS in the spinal cord and brain stem are considered in transitional zone (Fraher, 1992). The enlarged neurites are not found in the distal rootlets and DRGs in the aged animals. Compare to the other segments of the spinal cords, the megaloneurites occurred in the caudal lumber, sacral and rostral coccygeal in aged dogs. All these of segmental selective malformed structures demonstrate that the caudal of spinal cord especially in the sacral segments exit related adverse conditions more vulnerable to aging alterations.

Studies have shown that projections from the DGC have been also implicated in visceral nociception as discrete lesions of the dorsal column have been found to relieve pelvic pain in patients (Al-Chaer et al., 1996; Willis et al., 1999; Palecek and Willis, 2003; Liao et al., 2015). This clinical evidence is supported by anatomical studies using rats and monkeys. The axons of the neurons in the area adjacent to the CC including the ventral DGC, where nociceptive neurons are located, have been shown to travel in the dorsal column to the gracile nucleus and in the ventrolateral quadrant to the reticular formation (Al-Chaer et al., 1996; Wang et al., 1999; Liao et al., 2015).

Neuropathological studies have shown that NADPH-d staining could be used to reveal the neuronal terminal-pathy of aged conditions(Ma et al., 2000) as well as neurodegenerative animal models (Quinn et al., 2001). The experiments have demonstrated NADPH-d positive large-and medium-sized cells were multipolar in shape with dendrites and axon terminals together with a few dense fiber networks and evident in the gracile nucleus of aged rats (24 months old). We do not find significant NADPH-d neuritic alterations in the gracile nucleus and DRG in aged dogs. In our unpublished study in aged rats, the aging related NADPH-d neurodegeneration was detected in the gracile nucleus, which is similarly derived from the aging alterations in the lumbosacral spinal cord (Tan et al., 2006; Wei et al, 2016). The inputs of the primary sensory neurons of DRG could form axon bifurcation in the spinal cord. The bifurcating collaterals can terminal in the corresponding spinal segments and ascend to dorsal column nuclei respectively (Réthelyi and Szentágothai, 1973). Beside somatic sensory inputs, dorsal column nuclei also receive ascending visceral sensory information (Willis et al., 1999). The spheroidal NADPH-d neurodegenerations in the lumbosacarl spinal cord and gracile nucleus was postulated for dying back in aged rats (Blakemore and Cavanagh, 1969; Oda et al., 1992).

Different to the rats, concomitant with the prominent megaloneurites or the anomalous structures are found around the CC in the aged dogs. Tang et al report that NADPH-d positive nerve fibers formed a subependymal plexus and occurred to traverse the ependyma to run internally along the CC lumen (Tang et al., 1995). The NADPH-d ependymal traverse neurons could work as the cerebrospinal fluid (CSF)-contacting neurons. Bringing together our finding, aging NADPH-d neuritic hypertrophy around the CC was postulated as a neuritic dilation of neuronal fiber crossed ependyma, formed one kind of neurodegeneation. The CSF-contacting neurons function as pH sensors and mechanoreceptors (Jalalvand et al., 2018). Depleting the CSF-contacting neurons by neurotoxicity do not cause vital status(Song and Zhang, 2018). The megaloneurites and similarity around the CC may produce a localized disturbance and could then block axonal transport to give rise to morphological alterations identical to denervation.

The central projection of NADPH-d afferent fibers in the rat correspond to several of the sites in the lumbosacral spinal cord that have been shown to receive afferent input from the pelvic viscera. For example, measurements of increased expression of immediate early genes (e.g., c-fos) have revealed that nociceptive and nonnociceptive afferents from the urinary bladder(Birder et al., 1991; Birder and de Groat, 1992) and colon-rectum (Traub et al., 1992) activate cells in the region of the LCP and SPN corresponding to those sites exhibiting prominent NADPH-d fiber staining. In the rat, increased c-fos expression was induced in spinal tract neurons, interneurons, and parasympathetic preganglionic neurons (Ivanusic, 2008; Qi and Li, 2012). In the thoracic spinal cord, the number of NADPH-d positive neurons in the IML did not change. We did not find the NADPH-d positive neurons in the SPN, but the number of NADPH-d positive neurons in the DGC decreased in the aged dogs. The number of sympathetic neurons remained relative stable. We believe that the sympathetic neurons is important for vital organs. While the neurons innervating pelvic organs may be more vulnerable to aging deterioration.

NADPH-d activity in neurons and fibers in the DRG provides some support for the proposal that NADPH-d reactivity is present in pelvic visceral afferent pathways. NADPH-d staining in DRG cells occurred with different intensities ranging from intense to very light. Intensely stained NADPH-d DRG cells, like visceral afferent and dorsal root ganglion cells, are among the smallest cells in the ganglia (Kawatani et al., 1986; Vizzard et al., 1994a). This raises the possibility that the NADPH-d fiber bundle in the LCP might represent the central projections of the small cells. The pelvic visceral afferents enter the spinal dorsal horn at dorsal root entry zone and continue with LT. Many previous studies have shown that in patients with nerve injury, impaired sensory afferents enter the posterolateral aspect of the spinal cord through the dorsal root entry zone, and dorsal root entry zone lesions can alleviate neuropathic pain (Matt S. Ramer, 2001; Yang et al., 2015).

### Conclusion

NADPH-d neuronal positivity widely distributes in all segments of the spinal cord and brain. However, our finding is regarding selective neuronal vulnerability attributed to regional, cell-type and aging. Although the megaloneurites and similarity alterations selectively occurred in the sacral spinal cord of aged dogs, it is hardly determined that the subpopulations of NADPH-d neuronal structures are of segmental variation. The cellular senescence and aging neurodegeneration may determine the other intrinsic vulnerabilities of the sacral spinal cord(McHugh and Gil, 2018). We do not know if other biochemical property of similar enlarged fibers exited. The further experiments are required to prove the hypothesis. In general, we found two kind of aging changes in spinal cord of aged dogs, non-specific and specific segmental alteration. The number of NADPH-d neurons significantly reduces in entire spinal cord without clear segmental variation. It is an evidence for a pan-neurodegenerative change. However, the megaloneurites selectively occur in the caudal spinal cord of aged dogs, which implicated aging dysfunction of urogenital organs.

In summary, the major new finding revealed by this study is that the NADPH-d positive megaloneurites colocalize with VIP in the aged lumbosacral spinal cord, but not in young rats. The megaloneurites might be a swelling of the transganglionic fibers located in the dorsal root entry zoon, LCP in the lateral dorsal marginal to the dorsal horn, SPN, the DGC and the white matter as well as the CC in the sacral spinal cord. We think that the megaloneurites are considered as a specific aging marker, age-associated progressive deterioration and malformed structure.

## Acknowledgments

This work was supported by grants from National Natural Science Foundation of China (81471286), Liaoning Training Programs of scientific research and career development for Undergraduates, 201410160007 and Research Start-Up Grant for New Science Faculty of Jinzhou Medical University.

## References

Aimi Y, Fujimura M, Vincent SR, Kimura H. 1991. Localization of NADPH-diaphorase-containing neurons in sensory ganglia of the rat. The Journal of comparative neurology 306(3):382–392.

Al-Chaer ED, Lawand NB, Westlund KN, Willis WD. 1996. Visceral nociceptive input into the ventral posterolateral nucleus of the thalamus: a new function for the dorsal column pathway. J Neurophysiol 76(4):2661–2674.

Anderson CR. 1992. NADPH diaphorase-positive neurons in the rat spinal cord include a subpopulation of autonomic preganglionic neurons. Neuroscience letters 139(2):280–284.

Bansal U, Fuller TW, Jiang X, Bandari J, Zhang Z, Shen B, Wang J, Roppolo JR, de Groat WC, Tai C. 2017. Lumbosacral spinal segmental contributions to tibial and pudendal neuromodulation of bladder overactivity in cats. Neurourology and urodynamics 36(6):1496–1502.

Barbe MF, Gomez-Amaya SM, Salvadeo DM, Lamarre NS, Tiwari E, Cook S, Glair CP, Jang DH, Ragheb RM, Sheth A, Braverman AS, Ruggieri MR. 2018. Clarification of the Innervation of the Bladder, External Urethral Sphincter and Clitoris: A Neuronal Tracing Study in Female Mongrel Hound Dogs. Anatomical record (Hoboken, NJ : 2007).

Belai A, Schmidt HH, Hoyle CH, Hassall CJ, Saffrey MJ, Moss J, Forstermann U, Murad F, Burnstock G. 1992. Colocalization of nitric oxide synthase and NADPH-diaphorase in the myenteric plexus of the rat gut. Neuroscience letters 143(1-2):60–64.

Berkley KJ, Hubscher CH, Wall PD. 1993. Neuronal responses to stimulation of the cervix, uterus, colon, and skin in the rat spinal cord. J Neurophysiol 69(2):545–556.

Birder LA, de Groat WC. 1992. Increased c-fos expression in spinal neurons after irritation of the lower urinary tract in the rat. J Neurosci 12(12):4878–4889.

Birder LA, Roppolo JR, Iadarola MJ, de Groat WC. 1991. Electrical stimulation of visceral afferent pathways in the pelvic nerve increases c-fos in the rat lumbosacral spinal cord. Neuroscience letters 129(2):193–196.

Blakemore WF, Cavanagh JB. 1969. “Neuroaxonal dystrophy” occurring in an experimental “dying back” process in the rat. Brain 92(4):789–804.

Blok BF, Holstege G. 2000. The pontine micturition center in rat receives direct lumbosacral input. An ultrastructural study. Neuroscience letters 282(1-2):29–32.

Chakder S, Rattan S. 1996. Evidence for VIP-induced increase in NO production in myenteric neurons of opossum internal anal sphincter. The American journal of physiology 270(3 Pt 1):G492–497.

Chertok VM, Kotsuba AE. 2013. The distribution of NADPH-diaphorase and neuronal no synthase in rat medulla oblongata nuclei. Morfologiia (Saint Petersburg, Russia) 144(6):9–14.

Cohen BA, Major MR, Huizenga BA. 1991. Pudendal nerve evoked potential monitoring in procedures involving low sacral fixation. Spine (Phila Pa 1976) 16(8 Suppl):S375–378.

Cruz Y, Hernandez-Plata I, Lucio RA, Zempoalteca R, Castelan F, Martinez-Gomez M. 2017. Anatomical organization and somatic axonal components of the lumbosacral nerves in female rabbits. Neurourology and urodynamics 36(7):1749–1756.

Dawson TM, Bredt DS, Fotuhi M, Hwang PM, Snyder SH. 1991. Nitric oxide synthase and neuronal NADPH diaphorase are identical in brain and peripheral tissues. Proceedings of the National Academy of Sciences of the United States of America 88(17):7797–7801.

Doone VG, Pelissier N, Manchester T, Vizzard AM. 1999. Distribution of NADPH-d and nNOS-IR in the thoracolumbar and sacrococcygeal spinal cord of the guinea pig. Journal of the autonomic nervous system 77(2-3):98–113.

Fraher JP. 1992. The CNS-PNS transitional zone of the rat. Morphometric studies at cranial and spinal levels. Prog Neurobiol 38(3):261–316.

Gravholt CH, Naeraa RW, Fisker S, Christiansen JS. 1997. Body composition and physical fitness are major determinants of the growth hormone-insulin-like growth factor axis aberrations in adult Turner’s syndrome, with important modulations by treatment with 17 beta-estradiol. The Journal of clinical endocrinology and metabolism 82(8):2570–2577.

Hope BT, Michael GJ, Knigge KM, Vincent SR. 1991. Neuronal NADPH diaphorase is a nitric oxide synthase. Proceedings of the National Academy of Sciences of the United States of America 88(7):2811–2814.

Ivanusic JJ. 2008. The pattern of Fos expression in the spinal dorsal horn following acute noxious mechanical stimulation of bone. Eur J Pain 12(7):895–899.

Jalalvand E, Robertson B, Tostivint H, Löw P, Wallén P, Grillner S. 2018. Cerebrospinal fluid-contacting neurons sense pH changes and motion in the hypothalamus. The Journal of Neuroscience 38(35):7113–7724.

Jobling P, Graham BA, Brichta AM, Callister RJ. 2010. Cervix stimulation evokes predominantly subthreshold synaptic responses in mouse thoracolumbar and lumbosacral superficial dorsal horn neurons. J Sex Med 7(6):2068–2076.

Kawatani M, Nagel J, de Groat WC. 1986. Identification of neuropeptides in pelvic and pudendal nerve afferent pathways to the sacral spinal cord of the cat. The Journal of comparative neurology 249(1):117–132.

Kluchova D, Rybarova S, Schmidtova K, Kocisova M. 1999. Difference of pericentral NADPH-d positive neurons in the rabbit spinal cord segments. General physiology and biophysics 18 Suppl 1:69–71.

Kuo DC, de Groat WC. 1985. Primary afferent projections of the major splanchnic nerve to the spinal cord and gracile nucleus of the cat. The Journal of comparative neurology 231(4):421–434.

Liao CC, DiCarlo GE, Gharbawie OA, Qi HX, Kaas JH. 2015. Spinal cord neuron inputs to the cuneate nucleus that partially survive dorsal column lesions: A pathway that could contribute to recovery after spinal cord injury. The Journal of comparative neurology 523(14):2138–2160.

Liu Y, Tan H, Wan X, Zuo Z, Liu K. 1998. Spinal segment distribution of neural innervation related to houhai acupoint and compared with zusanli and dazhui acupoints in domestic chicken. Zhongguo Yi Xue Ke Xue Yuan Xue Bao 20(2):154–160.

Lynn RB, Sankey SL, Chakder S, Rattan S. 1995. Colocalization of NADPH-diaphorase staining and VIP immunoreactivity in neurons in opossum internal anal sphincter. Digestive diseases and sciences 40(4):781–791.

Ma S, Cornford ME, Vahabnezhad I, Wei S, Li X. 2000. Responses of nitric oxide synthase expression in the gracile nucleus to sciatic nerve injury in young and aged rats. Brain Res 855(1):124–131.

Marsala J, Marsala M, Lukacova N, Ishikawa T, Cizkova D. 2003. Localization and distribution patterns of nicotinamide adenine dinucleotide phosphate diaphorase exhibiting axons in the white matter of the spinal cord of the rabbit. Cell Mol Neurobiol 23(1):57–92.

Marsala J, Marsala M, Vanicky I, Taira Y. 1999. Localization of NADPHd-exhibiting neurons in the spinal cord of the rabbit. The Journal of comparative neurology 406(2):263–284.

Marsala J, Vanicky I, Marsala M, Jalc P, Orendacova J, Taira Y. 1998. Reduced nicotinamide adenine dinucleotide phosphate diaphorase in the spinal cord of dogs. Neuroscience 85(3):847–862.

Matsumoto T, Nakane M, Pollock JS, Kuk JE, Forstermann U. 1993. A correlation between soluble brain nitric oxide synthase and NADPH-diaphorase activity is only seen after exposure of the tissue to fixative. Neuroscience letters 155(1):61–64.

Matt S., Ramer ID. 2001. Two-Tiered Inhibition of Axon Regeneration at the Dorsal Root. The Journal of Neuroscience 21(8):2651–2660.

McHugh D, Gil J. 2018. Senescence and aging: Causes, consequences, and therapeutic avenues. The Journal of Cell Biology 217(1):65–77.

McKenna KE, Nadelhaft I. 1986. The organization of the pudendal nerve in the male and female rat. The Journal of comparative neurology 248(4):532–549.

McNeill DL, Traugh NE, Jr., Vaidya AM, Hua HT, Papka RE. 1992. Origin and distribution of NADPH-diaphorase-positive neurons and fibers innervating the urinary bladder of the rat. Neuroscience letters 147(1):33–36.

Morgan C, Nadelhaft I, de Groat WC. 1981. The distribution of visceral primary afferents from the pelvic nerve to Lissauer’s tract and the spinal gray matter and its relationship to the sacral parasympathetic nucleus. The Journal of comparative neurology 201(3):415–440.

Nadelhaft I, Booth AM. 1984. The location and morphology of preganglionic neurons and the distribution of visceral afferents from the rat pelvic nerve: a horseradish peroxidase study. The Journal of comparative neurology 226(2):238–245.

Nadelhaft I, Vera PL. 1996. Neurons in the rat brain and spinal cord labeled after pseudorabies virus injected into the external urethral sphincter. The Journal of comparative neurology 375(3):502–517.

O’Kelly TJ, Davies JR, Brading AF, Mortensen NJ. 1994. Distribution of nitric oxide synthase containing neurons in the rectal myenteric plexus and anal canal. Morphologic evidence that nitric oxide mediates the rectoanal inhibitory reflex. Dis Colon Rectum 37(4):350–357.

Oda K, Yamazaki K, Miura H, Shibasaki H, Kikuchi T. 1992. Dying back type axonal degeneration of sensory nerve terminals in muscle spindles of the gracile axonal dystrophy (GAD) mutant mouse. Neuropathol Appl Neurobiol 18(3):265–281.

Ogiwara H, Morota N. 2014. Pudendal afferents mapping in posterior sacral rhizotomies. Neurosurgery 74(2):171–175.

Orendacova J, Marsala M, Sulla I, Kafka J, Jalc P, Cizkova D, Taira Y, Marsala J. 2000. Incipient cauda equina syndrome as a model of somatovisceral pain in dogs: spinal cord structures involved as revealed by the expression of c-fos and NADPH diaphorase activity. Neuroscience 95(2):543–557.

Palecek J, Willis WD. 2003. The dorsal column pathway facilitates visceromotor responses to colorectal distention after colon inflammation in rats. Pain 104(3):501–507.

Porseva VV, Shilkin VV. 2010. NADPH-diaphorase-positive structures in the spinal cord and spinal ganglia. Morfologiia (Saint Petersburg, Russia) 137(2):13–17.

Portiansky EL, Barbeito CG, Gimeno EJ, Zuccolilli GO, Goya RG. 2006. Loss of NeuN immunoreactivity in rat spinal cord neurons during aging. Exp Neurol 202(2):519–521.

Pullen AH, Humphreys P. 1995. Diversity in localisation of nitric oxide synthase antigen and NADPH-diaphorase histochemical staining in sacral somatic motor neurones of the cat. Neuroscience letters 196(1-2):33–36.

Qi DB, Li WM. 2012. Effects of electroacupuncture on expression of c-fos protein in the spinal dorsal horn of rats with chronic visceral hyperalgesia. Zhong Xi Yi Jie He Xue Bao 10(12):1490–1496.

Qi HX, Kaas JH. 2006. Organization of primary afferent projections to the gracile nucleus of the dorsal column system of primates. The Journal of comparative neurology 499(2):183–217.

Quinn J, Davis F, Woodward WR, Eckenstein F. 2001. Beta-amyloid plaques induce neuritic dystrophy of nitric oxide-producing neurons in a transgenic mouse model of Alzheimer’s disease. Exp Neurol 168(2):203–212.

Réthelyi M, Szentágothai J. 1973. Distribution and Connections of Afferent Fibres in the Spinal Cord. In: Albe-Fessard D, Andres KH, Bates JAV, Besson JM, Brown AG, Burgess PR, Darian-Smith I, v. Düring M, Gordon G, Hensel H, Jones E, Libet B, Oscarsson O, Perl ER, Pompeiano O, Powell TPS, Réthelyi M, Schmidt RF, Semmes J, Skoglund S, Szentágothai J, Towe AL, Wall PD, Werner G, Whitsel BL, Zotterman Y, Iggo A, eds. Somatosensory System. Berlin, Heidelberg: Springer Berlin Heidelberg. p 207–252.

Saito S, Kidd GJ, Trapp BD, Dawson TM, Bredt DS, Wilson DA, Traystman RJ, Snyder SH, Hanley DF. 1994. Rat spinal cord neurons contain nitric oxide synthase. Neuroscience 59(2):447–456.

Santer RM, Dering MA, Ranson RN, Waboso HN, Watson AHD. 2002. Differential susceptibility to ageing of rat preganglionic neurones projecting to the major pelvic ganglion and of their afferent inputs. Autonomic Neuroscience 96(1):73–81.

Song SY, Zhang LC. 2018. The Establishment of a CSF-Contacting Nucleus “Knockout” Model Animal. Front Neuroanat 12:22.

Spike RC, Todd AJ, Johnston HM. 1993. Coexistence of NADPH diaphorase with GABA, glycine, and acetylcholine in rat spinal cord. The Journal of comparative neurology 335(3):320–333.

Steers WD, Ciambotti J, Etzel B, Erdman S, de Groat WC. 1991. Alterations in afferent pathways from the urinary bladder of the rat in response to partial urethral obstruction. The Journal of comparative neurology 310(3):401–410.

Tamura M, Kagawa S, Kimura K, Kawanishi Y, Tsuruo Y, Ishimura K. 1994. Distribution of NADPH diaphorase-positive nerves in human penile tissue. Nihon Hinyokika Gakkai zasshi The japanese journal of urology 85(11):1643–1648.

Tamura M, Kagawa S, Kimura K, Kawanishi Y, Tsuruo Y, Ishimura K. 1995. Coexistence of nitric oxide synthase, tyrosine hydroxylase and vasoactive intestinal polypeptide in human penile tissue--a triple histochemical and immunohistochemical study. The Journal of urology 153(2):530–534.

Tan H, He J, Wang S, Hirata K, Yang Z, Kuraoka A, Kawabuchi M. 2006. Age-related NADPH-diaphorase positive bodies in the lumbosacral spinal cord of aged rats. Archives of histology and cytology 69(5):297–310.

Tang FR, Tan CK, Ling EA. 1995. The distribution of NADPH-d in the central grey region (lamina X) of rat upper thoracic spinal cord. J Neurocytol 24(10):735–743.

Thor KB, Morgan C, Nadelhaft I, Houston M, De Groat WC. 1989. Organization of afferent and efferent pathways in the pudendal nerve of the female cat. The Journal of comparative neurology 288(2):263–279.

Traub RJ, Pechman P, Iadarola MJ, Gebhart GF. 1992. Fos-like proteins in the lumbosacral spinal cord following noxious and non-noxious colorectal distention in the rat. Pain 49(3):393–403.

Traub RJ, Solodkin A, Meller ST, Gebhart GF. 1994. Spinal cord NADPH-diaphorase histochemical staining but not nitric oxide synthase immunoreactivity increases following carrageenan-produced hindpaw inflammation in the rat. Brain Res 668(1-2):204–210.

Valtschanoff JG, Weinberg RJ, Rustioni A. 1992. NADPH diaphorase in the spinal cord of rats. The Journal of comparative neurology 321(2):209–222.

Verstegen AMJ, Vanderhorst V, Gray PA, Zeidel ML, Geerling JC. 2017. Barrington’s nucleus: Neuroanatomic landscape of the mouse “pontine micturition center”. The Journal of comparative neurology 525(10):2287–2309.

Vizzard MA, Erdman SL, de Groat WC. 1993a. The effect of rhizotomy on NADPH diaphorase staining in the lumbar spinal cord of the rat. Brain Res 607(1-2):349–353.

Vizzard MA, Erdman SL, de Groat WC. 1993b. Localization of NADPH-diaphorase in pelvic afferent and efferent pathways of the rat. Neuroscience letters 152(1-2):72–76.

Vizzard MA, Erdman SL, de Groat WC. 1993c. Localization of NADPH diaphorase in bladder afferent and postganglionic efferent neurons of the rat. Journal of the autonomic nervous system 44(1):85–90.

Vizzard MA, Erdman SL, de Groat WC. 1996. Increased expression of neuronal nitric oxide synthase in bladder afferent pathways following chronic bladder irritation. The Journal of comparative neurology 370(2):191–202.

Vizzard MA, Erdman SL, Erickson VL, Stewart RJ, Roppolo JR, De Groat WC. 1994a. Localization of NADPH diaphorase in the lumbosacral spinal cord and dorsal root ganglia of the cat. The Journal of comparative neurology 339(1):62–75.

Vizzard MA, Erdman SL, Forstermann U, de Groat WC. 1994b. Differential distribution of nitric oxide synthase in neural pathways to the urogenital organs (urethra, penis, urinary bladder) of the rat. Brain Res 646(2):279–291.

Vizzard MA, Erdman SL, Roppolo JR, Forstermann U, de Groat WC. 1994c. Differential localization of neuronal nitric oxide synthase immunoreactivity and NADPH-diaphorase activity in the cat spinal cord. Cell Tissue Res 278(2):299–309.

Vizzard MA, Erickson K, de Groat WC. 1997. Localization of NADPH diaphorase in the thoracolumbar and sacrococcygeal spinal cord of the dog. Journal of the autonomic nervous system 64(2-3):128–142.

Vizzard MA, Erickson VL, Card JP, Roppolo JR, de Groat WC. 1995. Transneuronal labeling of neurons in the adult rat brainstem and spinal cord after injection of pseudorabies virus into the urethra. The Journal of comparative neurology 355(4):629–640.

Wang CC, Willis WD, Westlund KN. 1999. Ascending projections from the area around the spinal cord central canal: A Phaseolus vulgaris leucoagglutinin study in rats. The Journal of comparative neurology 415(3):341–367.

Wang HF, Shortland P, Park MJ, Grant G. 1998. Retrograde and transganglionic transport of horseradish peroxidase-conjugated cholera toxin B subunit, wheatgerm agglutinin and isolectin B4 from Griffonia simplicifolia I in primary afferent neurons innervating the rat urinary bladder. Neuroscience 87(1):275–288.

Willis WD, Al-Chaer ED, Quast MJ, Westlund KN. 1999. A visceral pain pathway in the dorsal column of the spinal cord. Proceedings of the National Academy of Sciences of the United States of America 96(14):7675–7679.

Wolf HK, Buslei R, Schmidt-Kastner R, Schmidt-Kastner PK, Pietsch T, Wiestler OD, Blumcke I. 1996. NeuN: a useful neuronal marker for diagnostic histopathology. J Histochem Cytochem 44(10):1167–1171.

Yang F, Zhang C, Xu Q, Tiwari V, He SQ, Wang Y, Dong X, Vera-Portocarrero LP, Wacnik PW, Raja SN, Guan Y. 2015. Electrical stimulation of dorsal root entry zone attenuates wide-dynamic-range neuronal activity in rats. Neuromodulation 18(1):33–40; discussion 40.

Zhou Y, Ling EA. 1998. Colocalization of nitric oxide synthase and some neurotransmitters in the intramural ganglia of the guinea pig urinary bladder. The Journal of comparative neurology 394(4):496–505.

Zhou Y, Tan CK, Ling EA. 1997. Distribution of NADPH-diaphorase and nitric oxide synthase-containing neurons in the intramural ganglia of guinea pig urinary bladder. Journal of anatomy 190 (Pt 1):135–145.

Wei Z, Xu X, Wang Y, Wen X, Li F, Shu G, Huang Y, Wu X, Li H, Rao C, Sun J, Wu W, Jiang H, Bai L, Li Y, Zheng D, Li D, Jiang D, Shi G, Du G, Zhai Z, Tan H. 2016. Aging-related NADPH-diaphorase Body (Tan body): a Morphological Marker of Aging Neurodegenerative Damage.

